# Pharmacologic inhibition of OLIG2 by CT-179 disrupts mitotic progression and enhances standard-of-care chemoradiotherapy in glioblastoma

**DOI:** 10.64898/2026.10.14.759257

**Authors:** Sam A. Greenall, Shengnan C. Chen, Gordon Alton, Graham Beaton, Reghan Boelter, Mattie Casey, Ivan Babic, Zara C. Bruce, Daniel J. Garama, Rajesh Mukthavaram, Fabio C. Tucci, Satheesh B. Ravula, Ambrosio Lopez, Susan Knowles, Gregory D. Stein, Yuchen Li, Bennett Stork, Rochelle D’Souza, Ashwani Kumar, Ralf Kittler, Ann C.M. Tuma, Jann N. Sarkaria, Daniel J. Gough, Rodney A. Stewart, Santosh Kesari, Bryan W. Day, Terrance G. Johns

**Author notes:** Corresponding author: Terrance G. Johns, PhD. Latrobe University, Bundoora, Vic, Australia. Present addresses: Sam A. Greenall: Department of Medicine, Monash University, Clayton, VIC, Australia Gordon Alton: Agragene, Inc., San Diego, CA, USA. Graham Beaton: Epigen Biosciences, 10225 Barnes Canyon Road, Suite A104, San Diego, CA 92121, USA. Shengnan C. Chen: Medpace, Melbourne, VIC, Australia Reghan Boelter: Homespire Health, Salt Lake City, UT, USA Ivan Babic: CellarisBio, San Diego, CA, USA. Rajesh Mukthavaram: Arcturus Therapeutics Ltd, San Diego, CA, USA Susan Knowles: CD Bradshaw & Associates, Georgetown, TX, USA Terrance G. Johns: Latrobe University, Bundoora, Vic, Australia. Santosh Kesari: Asthra Health, Santa Monica, CA, and Lundquist Institute, Torrance, CA, USA. Sam A. Greenall and Shengnan C. Chen contributed equally to this work and are designated joint first authors.

## Abstract

**Background:** Glioblastoma (GBM) remains uniformly fatal. The basic helix-loop-helix transcription factor OLIG2 is universally expressed in diffuse glioma, sustains the tumor-propagating stem-like fraction, and opposes p53-mediated genotoxic responses, but has historically been considered undruggable. We characterized CT-179, a brain-penetrant small-molecule OLIG2 inhibitor, as a candidate therapeutic for GBM.

**Methods:** Target engagement was assessed by cellular thermal shift assay, differential scanning fluorimetry, crosslinking, and electrophoretic mobility shift assay. Anti-tumor activity was evaluated across eleven patient-derived GBM cell lines, in zebrafish oligodendrocyte precursor cell (OPC)-derived tumors, and orthotopic and subcutaneous GBM models, including combinations with radiotherapy plus temozolomide and with selective Bcl-2 family inhibitors. **Results:** CT-179 engaged OLIG2 and altered the conformation of dimerized OLIG2 on chromatin without preventing dimerization or fully displacing OLIG2 from DNA. CT-179 reduced OLIG2-positive OPC’s and OPC-derived tumors in zebrafish and inhibited proliferation across all eleven GBM lines (IC50 < 360 nM), depleting the stem-like fraction and inducing apoptosis through mitotic catastrophe, mitotic slippage, or a hybrid response. RNA-seq across three patient-derived glioma stem cell lines identified β-tubulin gene isoforms as the most strongly CT-179-down-regulated transcripts at the early time point and showed correlated β-tubulin and OLIG2 expression in TCGA glioblastoma specimens. Combination with the Mcl-1 inhibitor S63845 or the Bcl-2/Bcl-xL/Bcl-w inhibitor ABT-737 produced greater-than-additive cytotoxicity. Oral CT-179 crossed the blood-brain barrier and extended median survival as a single agent across two independent adult orthotopic patient-derived backgrounds (GBM8 and G06); intraperitoneal CT-179 was associated with reduced intratumoral OLIG2 and Ki-67 staining in GBM8 orthotopic xenografts. Combination of CT-179 with radiotherapy and temozolomide eliminated all GBM8 and 7 of 8 GBM4 subcutaneous xenografts without weight loss and significantly extended survival when combined with radiotherapy in MGMT-unmethylated GBM6 orthotopic xenografts (CT-179 administered intraperitoneally).

**Conclusions:** CT-179 is a brain-penetrant inhibitor of an established GBM driver with single-agent and combination activity across several orthotopic models, supporting clinical development for GBM and translational evaluation in other OLIG2-dependent CNS malignancies.

**Key Points:**

- CT-179 is the first brain-penetrant small-molecule inhibitor of OLIG2 with potent activity across patient-derived glioblastoma models.
- CT-179 induces apoptosis through mitotic catastrophe and slippage rather than collapse of receptor tyrosine kinase signaling.
- Oral CT-179 combined with radiotherapy and temozolomide eliminates the great majority of subcutaneous patient-derived GBM xenografts without overt toxicity.
- Oral CT-179 extends single-agent survival across two independent adult orthotopic patient-derived backgrounds (GBM8 and G06); intraperitoneal CT-179 is associated with reduced intratumoral OLIG2 and Ki-67 staining in GBM8 tumors.
- Intraperitoneal CT-179 plus radiotherapy significantly extends survival in an MGMT-unmethylated GBM6 orthotopic model where temozolomide contributes minimally and supports the rationale for further evaluation in MGMT-unmethylated GBM.

**Importance of the Study:** Glioblastoma is uniformly fatal, and durable disease-modifying therapies that disable the lineage transcription factor programs sustaining the GBM stem-like state have remained elusive. OLIG2 is universally expressed in diffuse glioma and is a central driver of stemness, lineage-restricted RTK signaling, and resistance to genotoxic therapy, yet has historically been considered undruggable. This study reports the foundational preclinical characterization of CT-179, a brain-penetrant small molecule that engages OLIG2 directly and induces glioblastoma cell death through disrupted mitotic progression. Oral CT-179 crosses the blood-brain barrier, prolongs survival in orthotopic patient-derived xenografts, and substantially enhances the efficacy of standard-of-care radiotherapy and temozolomide. Activity in an MGMT-unmethylated orthotopic patient-derived GBM model in combination with radiotherapy is of particular translational interest, since this is the patient population for which temozolomide contributes minimally and the unmet need is greatest. These data establish proof of concept for direct pharmacologic inhibition of OLIG2 in GBM and provide a translational rationale, including an MGMT-stratified combination strategy, for combining CT-179 with current standard-of-care chemoradiotherapy.

**Lay Summary:** Glioblastoma is the most common and aggressive brain cancer in adults, and almost all patients die of their disease. One reason is a protein called OLIG2, which keeps tumor cells in an immature, treatment-resistant state and has long been considered impossible to target with drugs. We studied CT-179, a small molecule that enters the brain and blocks OLIG2. In glioblastoma cells and in mice, CT-179 forced dividing tumor cells into a faulty, lethal division, extended survival on its own, and strengthened the effect of standard radiation and chemotherapy. These results support testing CT-179 in people with glioblastoma.

## Introduction

Glioblastoma (GBM), the most common malignant primary brain tumor in adults, remains uniformly fatal despite maximal surgical resection followed by radiotherapy and temozolomide (TMZ)^1,2^. Median overall survival has remained at 14–16 months for nearly two decades, and the 2021 World Health Organization (WHO) classification of central nervous system tumors has further consolidated GBM as a molecularly defined IDH-wildtype, grade 4 astrocytoma in which intratumoral heterogeneity, treatment-resistant stem-like populations, and an unfavorable molecular landscape together limit the durability of standard therapy^3,4^. New molecularly targeted approaches that disable the regulatory programs sustaining the GBM stem-like state are therefore needed.

Among the lineage transcription factors expressed in diffuse glioma, oligodendrocyte transcription factor 2 (OLIG2) occupies a central position. OLIG2 is a basic helix-loop-helix (bHLH) transcription factor that is universally expressed in diffuse gliomas across grade and histological subtype, and is largely restricted in the adult central nervous system to oligodendroglia and their progenitors, providing a favorable therapeutic window^5^. Genetic studies have established that OLIG2 is required for glioma formation in murine models, where it sustains the replication competence of neural stem and progenitor cells in part through direct repression of the cyclin-dependent kinase inhibitor p21^6^. OLIG2 also opposes p53-dependent responses to genotoxic stress, providing a mechanistic explanation for the relative resistance of OLIG2-positive gliomas to radiation and TMZ even when TP53 is structurally intact^7^. Beyond cell-cycle and stress-response control, OLIG2 marks and sustains the tumor-propagating, stem-like fraction of GBM and orchestrates a lineage-restricted shift in receptor tyrosine kinase (RTK) signaling that defines GBM molecular subtype^8–10^. Together, these data identify OLIG2 as both a marker and a driver of GBM and motivate its direct pharmacologic inhibition.

OLIG2 has historically been considered undruggable, in keeping with the broader challenge of inhibiting transcription factors. One emerging strategy for such targets is targeted protein degradation, in which heterobifunctional degraders (PROTACs) or molecular glues redirect an E3 ubiquitin ligase to deplete the protein of interest, an approach that has been proposed for glioma transcriptional regulators including OLIG2^11^. Computational work has, however, identified the OLIG2 dimerization interface as a tractable target. An in silico pharmacophore model of the OLIG2 bHLH domain enabled a virtual screen that yielded the first small-molecule OLIG2 inhibitors and provided proof of principle that the dimerization interface can be engaged by drug-like compounds^12,13^. Building on the screening hit CT-102, medicinal chemistry optimization has produced CT-179, a brain-penetrant urea-pyrimidine small molecule designed to engage OLIG2 within tumor cells^14,15^. In a parallel program, CT-179 has been shown to disrupt OLIG2 function in Sonic Hedgehog–subgroup medulloblastoma, where it suppresses OLIG2-positive tumor stem cells, increases differentiation, and potentiates radiotherapy in patient-derived xenograft and genetically engineered mouse models, but had no effect on normal cells^16^. These findings indicate that pharmacologic inhibition of OLIG2 is feasible and biologically active in at least one OLIG2-dependent malignancy, but the preclinical activity of CT-179 in adult GBM, the disease in which OLIG2 expression is most pervasive and for which a brain-penetrant OLIG2 inhibitor was originally conceived, has not been systematically defined. Here we characterize CT-179 as a small-molecule OLIG2 inhibitor for GBM. We demonstrate cellular target engagement, dissect the consequences of OLIG2 inhibition for oligodendrocyte precursor cell (OPC) biology in vivo and for the proliferation, stem-like fraction, RTK expression, and translational output of patient-derived GBM lines, and define the mitotic mechanisms through which CT-179 induces cell death. We further assess pharmacokinetic suitability for CNS indications and evaluate single-agent and combination activity in zebrafish and orthotopic murine models, including combination with radiotherapy and TMZ. The data position CT-179 as a brain-penetrant inhibitor of an established GBM driver and as a candidate for combination with current standard of care.

## Results

### CT-179 engages OLIG2 and functionally inhibits its DNA-binding activity

Building on a previously described in silico model of the OLIG2 dimerization interface^13^, we conducted a virtual screen that identified the hit compound CT-102 and, after extensive medicinal chemistry optimization, generated 1-{2-[3-(3,4-dichlorophenyl)ureido]-6-methyl-pyrimidin-4-yl}-amino-3-(dimethylamino)propane, designated CT-179^15^ (Fig. 1a). CT-179 was designed with physicochemical properties favorable for blood-brain barrier penetration. To confirm cellular target engagement, we performed cellular thermal shift assays (CETSA)^17,18^ in RR2 cells and observed enhanced OLIG2 thermal stability in the presence of CT-179 across a 40–50°C temperature gradient (Fig. 1b), consistent with direct binding.

**Figure 1:**
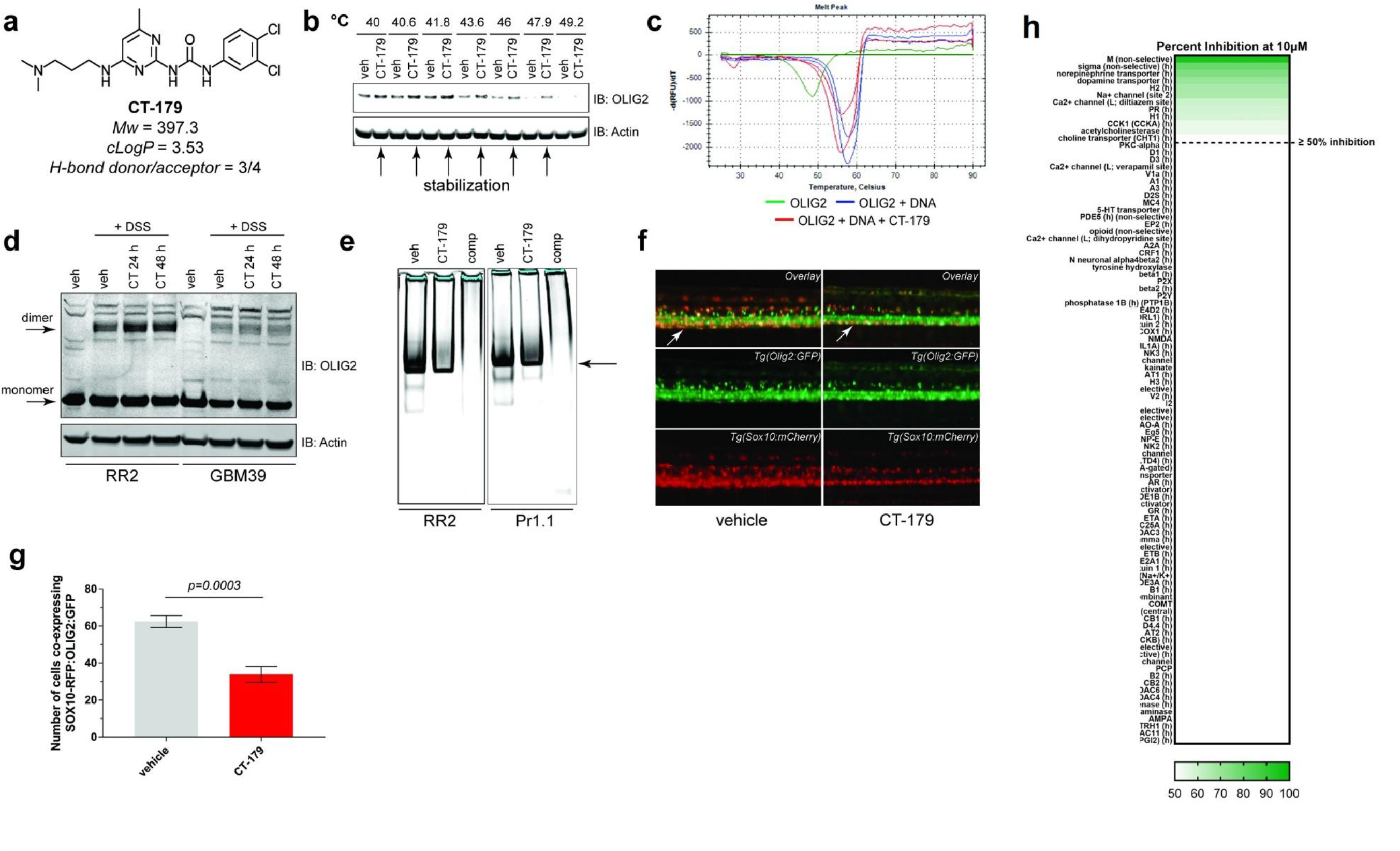
Validation of CT-179•OLIG2 engagement and effects of CT-179 on OLIG2 dimerization, DNA binding and in vivo transcriptional activity. a. Chemical structure and properties of CT-179. b. Western blot for OLIG2 status in RR2 cells after CETSA, using a temperature gradient between 40°C and 50°C. OLIG2 is stabilized (arrows) vs temperature matched vehicle controls in the presence of 20 µM CT-179 over several increasing temperatures. Actin was used as a loading control. c. Thermal shift assays on monomeric CT6-OLIG2 variant alone (green), CT6-OLIG2 bound in dimeric form to a MycN E-box dsDNA probe (blue) and CT6-OLIG2, dsDNA probe and CT-179 (red), which shows destabilization of the complex through a 1.8°C reduction in stability. d. Western blot for dimeric OLIG2 status following DSS crosslinking of RR2 and GBM39 cells after vehicle or 1 µM CT-179 treatment for the indicated times. Actin was used as a loading control. A non-crosslinked sample was included as a non-dimerized control. e. RR2 and Pr1.1 EMSA profiles for binding of endogenous OLIG2 within nuclear extracts to an IRDye800-labeled p27kip dsDNA probe in the presence or absence of 20 µM CT-179. An unlabeled p27^kip^ dsDNA competitor control (comp; 1000-fold excess) prevented the detected shift, confirming specificity. The arrow indicates the OLIG2•dsDNA p27^kip^ probe complex. f. Lateral views of the spinal cord of 5 dpf zebrafish embryos containing the Tg(*Olig2:eGFP*); Tg(*Sox10:mCherry*) reporters that simultaneously label developing OPCs (double labeling - white arrows) and differentiated oligodendrocytes (green only). Treatment with 15 µM CT-179 (right panels) starting at 2 dpf (48 hpf) for 3 days shows a clear reduction in the number of double-labeled OPCs (compare white arrows). **g.** Quantitation of images in **f.** Verification that CT-179 significantly reduces OPC numbers in vivo (62 cells vs 32 cells; n = 10, *P* = 0.0003 by unpaired two-tailed student’s t-test). **h.** Heat map of CT-179 inhibition across a 110-target binding and enzyme/uptake panel screened at 10 µM (Eurofins Cerep, Study #100022981). Targets are ordered by % inhibition; only 12 targets exceeded 50% inhibition, and only the muscarinic (M, non-selective) receptor exceeded 90% (94.4%). Reference compound performance was within historical specifications for each assay.

To probe the functional consequences of binding, we examined a recombinant bHLH-domain construct of OLIG2 (“CT6”) by differential scanning fluorimetry. Addition of CT-179 to preformed OLIG2•dsDNA complexes containing an E-box motif did not displace the protein from DNA but produced a reproducible 1.8°C decrease in the thermal stability of the dimerized OLIG2•dsDNA complex (Fig. 1c). Disuccinimidyl suberate (DSS) crosslinking in RR2 and GBM39 cells confirmed that CT-179 did not reduce the level of OLIG2 dimer (Fig. 1d).

Electrophoretic mobility shift assays (EMSA) in RR2 and Pr1.1 nuclear extracts further demonstrated that CT-179 reduced, but did not abolish, OLIG2 binding to an IRDye800-labeled p27kip E-box probe^19^ (Fig. 1e). Together, these data indicate that CT-179 engages an OLIG2• dsDNA complex and alters its conformation without preventing dimerization or fully displacing OLIG2 from DNA, suggesting that CT-179 modulates OLIG2 transcriptional activity through a conformational mechanism on chromatin. In a 110-target receptor, channel, and transporter panel screened at 10 µM (Eurofins Cerep), 12 targets exceeded 50% inhibition and only the muscarinic receptor exceeded 90% (Fig. 1h); kinome selectivity is described separately^16^. Because this panel was screened at 10 µM, well above the nanomolar concentrations that inhibit GBM cell proliferation and the brain exposures achieved in vivo, these secondary activities are unlikely at efficacious doses; the muscarinic activity nonetheless warrants monitoring for potential anticholinergic and central nervous system effects.

### CT-179 reduces OLIG2-positive oligodendrocyte precursor cell proliferation in vivo

Zebrafish provide a tractable system for evaluating the effect of OLIG2 inhibition on oligodendrocyte development. OPCs are multipotent CNS cells that give rise to myelinating oligodendrocytes; in zebrafish embryos, OPCs co-express the Olig2 and Sox10 transcription factors in the brain and spinal cord^20^. OPCs have also been implicated as a cell of origin for a subset of gliomas and embryonal brain tumors in mouse and zebrafish models^21,22^, motivating their use as a developmental readout of CT-179 activity.

We treated double-transgenic *Tg(Olig2:eGFP); Tg(Sox10:mCherry)* embryos at 2 days post-fertilization (dpf) with vehicle or 15 µM CT-179 for 3 days and quantified double-positive OPCs in the ventral spinal cord at 5 dpf. CT-179 treatment reduced OPC numbers by approximately 50% relative to vehicle controls (62 vs. 32 cells; n = 10, *P* = 0.0003; Fig. 1f,g and 1g), with no appreciable effect on differentiated GFP single-positive oligodendrocytes in the dorsal spinal cord. These observations indicate that CT-179 selectively reduces OLIG2-positive OPCs in a whole-animal model without overtly disrupting oligodendrocyte maturation. The micromolar bath concentrations used in these whole-animal immersion assays reflect the limited uptake of compound from the surrounding medium and are not directly comparable to the nanomolar IC50 values obtained in cell-based assays.

### CT-179 potently inhibits patient-derived GBM cell line proliferation

We screened CT-179 for anti-proliferative activity across eleven genetically and phenotypically diverse patient-derived GBM cell lines^23,24^. All cell lines were sensitive to CT-179, with IC50 values below 360 nM (Fig. 2a). Treatment with 1 µM CT-179 increased cell-surface Annexin V (Fig. 2b) and cleaved caspase 3/7 (Fig. 2c) across multiple lines, indicating that OLIG2 inhibition induces apoptosis. The kinetics of apoptosis induction varied between cell lines, with detectable responses ranging from 3 to 10 days post-treatment.

**Figure 2:**
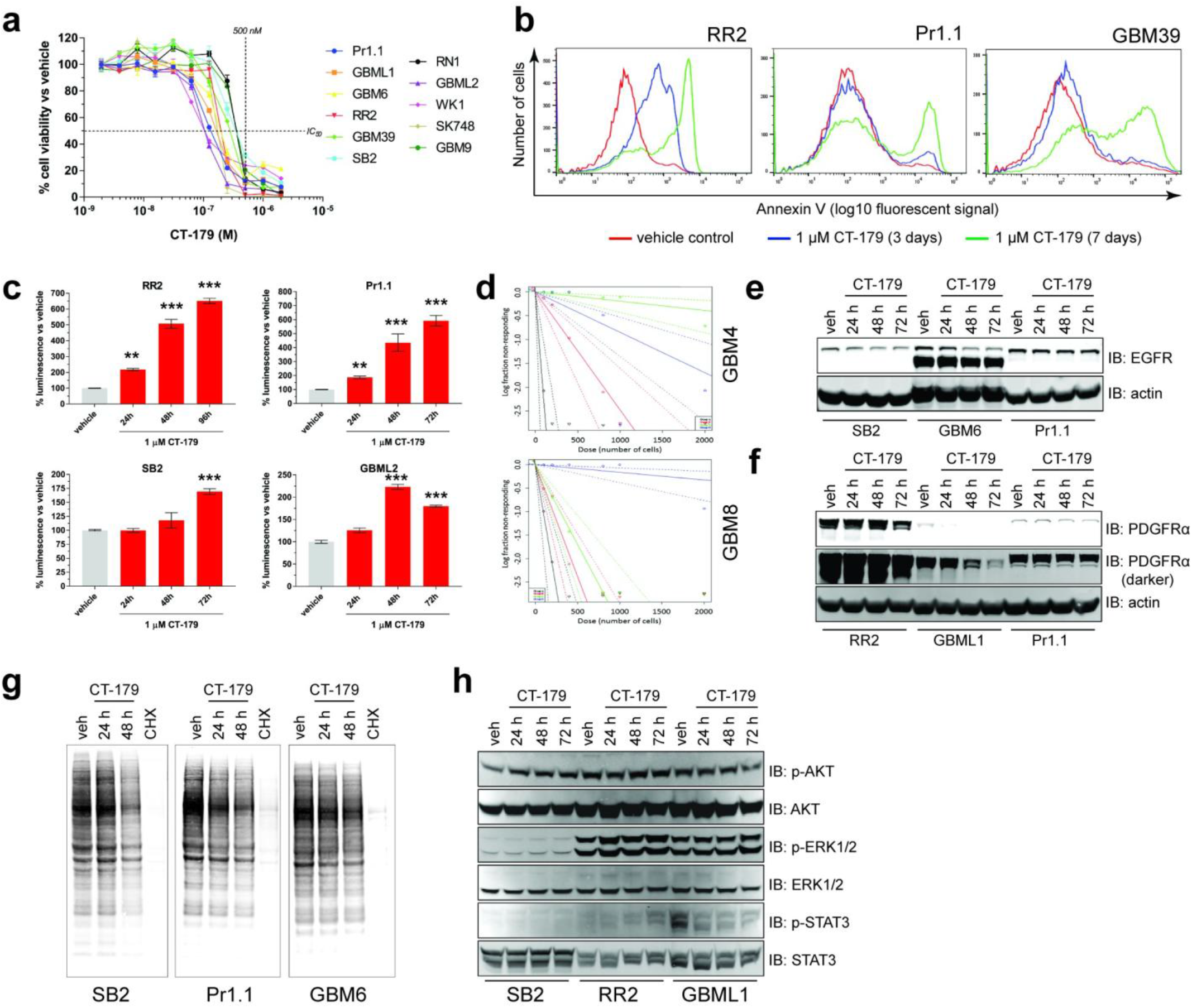
CT-179 treatment in several genotypically and phenotypically diverse primary GBM lines induces apoptosis and reduced global translation. a. Representative plots for the inhibition of proliferation in 11 primary GBM cell lines after 7 days of treatment with increasing concentrations of CT-179 as measured by Vialight assay. Data are presented as percentage of cell viability vs vehicle controls ± s.e.m. Horizontal dashed line = IC50 demarcation, vertical dashed line = 500 nM CT-179 demarcation. b. Flow cytometry histograms depicting cell surface Annexin V status of RR2, Pr1.1 and GBM39 cells treated with vehicle (red) or 1 µM CT-179 for 3 days (blue) or 7 days (green). c. Cleaved caspase 3/7 detection in RR2, Pr1.1, SB2 and GBML2 cells treated with 1 µM CT-179 for 24, 48 and 72 h. Data represent the percentage luminescence signal obtained for each time point vs vehicle controls ± s.e.m. ** represents *P* < 0.01, *** represents *P* < 0.001 significant difference vs vehicle controls by one-way ANOVA (n = 3 per test). **d.** Extreme limiting dilution plots for GBM4 and GBM8 following treatment with vehicle or various concentrations of CT-179. Black = vehicle. Red = 80 nM (GBM4) or 50 nM (GBM8). Blue = 250 nM (GBM4) or 75 nM (GBM8). Green = 750 nM (GBM4) or 100 nM (GBM8). **e-f.** Western blot analysis of EGFR/EGFRvIII (**e**) and PDGFR-α (**f**) expression following treatment of cells with vehicle or 1 µM CT-179 for the indicated times. Actin was used as a loading control. **g.** Nascent global translation product 35S labeling in the presence of vehicle, 1 µM CT-179 for 24 h or 48 h or 10 µM cycloheximide for 1 h in RR2, SB2, Pr1.1 and GBM6 cells. **h.** Western blot analysis of the activation status of three major RTK-downstream oncogenic signaling pathways (p-AKT, p-ERK1/2 and p-STAT3) in SB2, RR2 and GBML1 cell lines in response to treatment with vehicle or 1 µM CT-179 for the indicated times. Actin was used as a loading control.

A reduction in OLIG2 protein was observed in many cell lines following CT-179 treatment, occurring approximately 24 h after the onset of caspase cleavage and Annexin V positivity. This decrease was not attributable to loss of Olig2 transcription, and several cell lines (e.g., SB2 and GBM6) showed no reduction in OLIG2 protein despite undergoing apoptosis. Together, these data indicate that the decrease in OLIG2 levels is not required for the initiation of CT-179– mediated apoptosis but may contribute to the late stages of cell death.

### CT-179 dose-dependently reduces GBM stem cell ratios and affects RTK expression

OLIG2 is implicated in the maintenance of GBM stem-like populations and in the expression of receptor tyrosine kinases (RTKs) such as EGFR and PDGFR-α that contribute to gliomagenesis^25,26^. Extreme limiting dilution assays in GBM4 and GBM8 cells revealed a dose-dependent reduction in the ratio of GBM stem-like cells to total GBM cells across increasing concentrations of CT-179 (Fig. 2d), consistent with selective depletion of the stem-like compartment.

Western blot analyses showed that CT-179 reduced the protein levels of EGFR, PDGFR-α, and HER4, while expression of the EGFRvIII truncation mutant, MET, and HER2 was unaffected (Fig. 2e,f and 2f). The down-regulation of RTKs was not transcriptionally mediated, as *Egfr*, *Pdgfra*, *Met*, and *Erbb4* mRNA levels were unchanged, suggesting a translational mechanism. Consistent with this interpretation, pulse labeling of nascent protein demonstrated that CT-179 reduces global translation output (Fig. 2g) at time points coincident with RTK protein loss.

Despite the reduction in RTK expression, activation of ERK1/2, AKT, and STAT3 was sustained across six CT-179–treated cell lines (Fig. 2h), indicating that redundant or compensatory signaling inputs maintain downstream pathway activity in the face of partial RTK depletion.

### CT-179 disrupts mitotic progression to induce mitotic catastrophe in GBM cell lines

The persistence of activated RTK signaling in CT-179–treated cells indicates that RTK down-regulation alone does not account for the anti-tumor activity of CT-179, prompting us to examine cell-cycle effects. OLIG2 regulates the transcription of multiple cell-cycle effectors, and prior ChIP-seq studies indicate enrichment of OLIG2 target genes in pathways governing cell-cycle progression and microtubule function^27^, both of which converge on mitotic control. To define the proximate transcriptional consequences of OLIG2 inhibition, we performed RNA-seq on three patient-derived glioma stem cell lines (C419, CD133, and GBM4) at early (8–12 h) and late (36 h) time points after CT-179 treatment (36 libraries; FDR < 0.05). At the early time point, β-tubulin gene isoforms were among the most strongly down-regulated transcripts, with gene-set enrichment analysis (GSEA) yielding normalized enrichment scores of approximately −3 for the tubulin gene set in C419 cells (Supplementary Figure S3). At 36 h, expression changes were concordant across C419 and CD133, with shared up-regulated genes enriched for G2/M-phase transcripts and shared down-regulated genes enriched for G1/S-phase transcripts, a profile highly consistent with cells arresting in mitosis (Supplementary Figure S1). Critically, in TCGA glioblastoma specimens the three β-tubulin genes most strongly down-regulated by CT-179 (*TUBB2A*, *TUBB2B*, and *TUBB*) co-varied with OLIG2 expression at Spearman ρ = 0.551, 0.579, and 0.401, respectively (FDR-significant), nominating these β-tubulin isoforms as putative direct OLIG2 transcriptional targets in patient tumors. These analyses, however, rest on post-treatment transcript down-regulation and bulk-tumor co-expression rather than on demonstrated promoter occupancy; OLIG2 binding at the β-tubulin loci was not assessed here, and their direct regulation by OLIG2 therefore remains to be confirmed, for example by ChIP-seq. Cell-cycle analysis of asynchronous GBM cells treated with 1 µM CT-179 revealed a modest to profound G2/M-phase arrest in some lines and minimal arrest in others (Fig. 3a).

**Figure 3:**
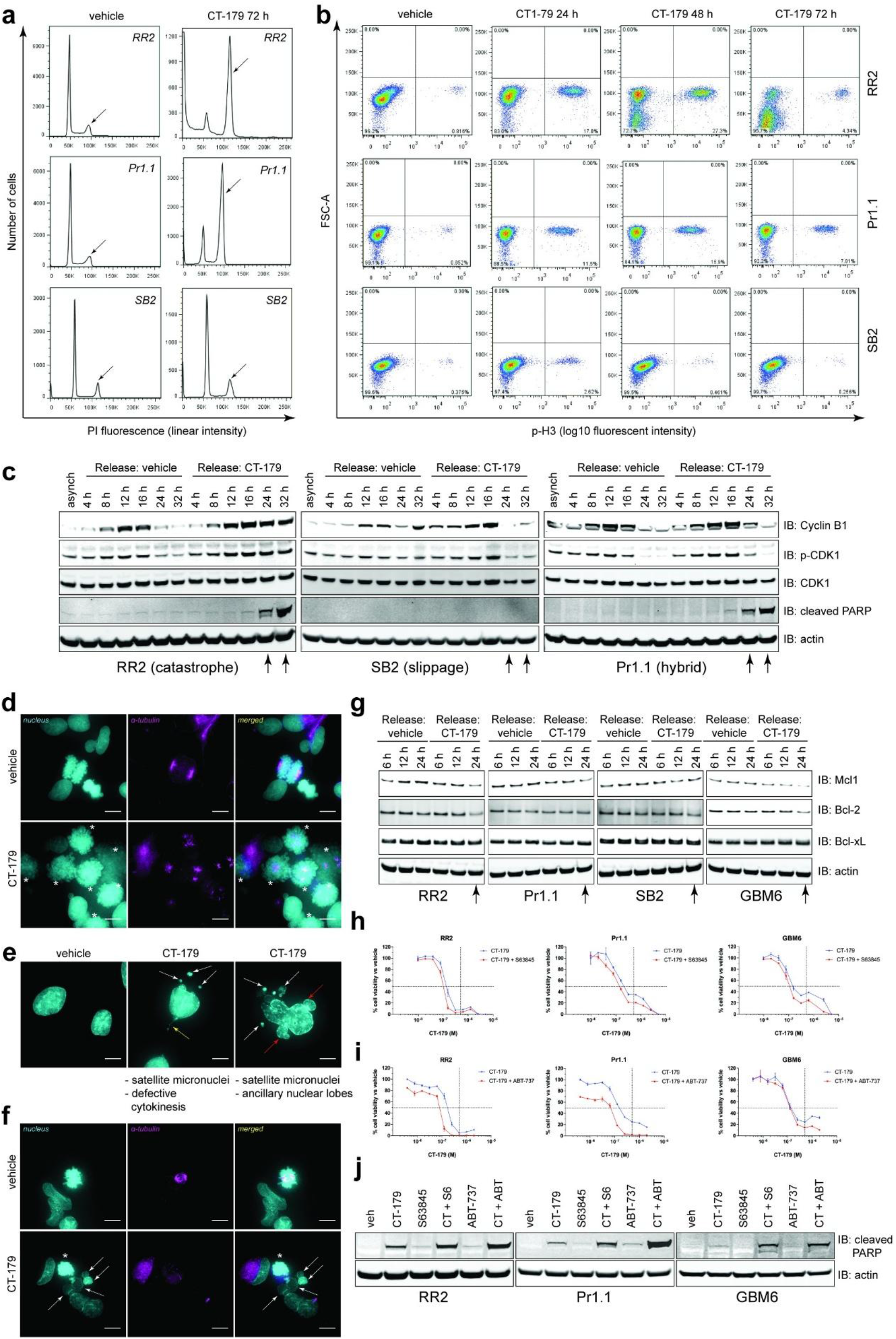
CT-179 disrupts mitotic fidelity, leading to cell death via mitotic catastrophe and/or mitotic slippage, resulting in apoptosis that can be enhanced by BH3 mimetics. **a.** Histograms of asynchronous RR2, Pr1.1 and SB2 cells fixed and stained with propidium iodide after 72 h treatment with vehicle (left panels) or 1 µM CT-179 (right panels) to establish the proportion of the total cell population in each cell cycle phase. The G2-M phase is depicted by the arrows. **b.** Flow cytometry plots depicting the pSer10-Histone 3 (p-H3) status of asynchronous RR2, Pr1.1 and SB2 cells treated with vehicle or 1 µM CT-179 for the indicated times. **c.** Western blot analyses of the major mitotic-phase proteins Cyclin B1, pThr161 and total CDK1 and the apoptotic protein cleaved PARP in RR2, Pr.1.1 and SB2 cells synchronized by double thymidine block and released from the G1-S boundary into vehicle or 1 µM CT-179 for the indicated time points. An asynchronous control (asynch) was included as well as an actin loading control. Arrows indicate the time points in CT-179 treated cells after which a full exit from the mitotic cycle is observed in vehicle controls. **d-f.** Immunofluorescence interrogation of the nuclear morphology and mitotic spindle status of RR2, SB2 and Pr1.1 cells treated with vehicle or 1 µM CT-179 for 48 h. Nucleus is stained by DAPI (cyan) and mitotic spindles are stained using anti-α-tubulin antibody (magenta). Scale bar = 10 µm. **d.** RR2 cells. Top panels demonstrate an anaphase cell with normal nuclear morphology and spindle alignment. Bottom panels show cells stalled at prometaphase with mitotic spindles formed but improperly aligned (asterisk), indicating mitotic catastrophe. **e.** SB2 cells. CT-179 treatment leads to abnormal nuclear phenotypes including residual cytokinetic bridges (yellow arrow), satellite micronuclei (white arrow) and/or ancillary nuclear lobe formation (red arrow), indicative of mitotic slippage. **f.** Pr1.1 cells. Top panels demonstrate an anaphase cell with normal nuclear morphology and spindle alignment. Bottom panels show a cell stalled at prometaphase with mitotic spindles formed but improperly aligned (asterisk) and a cell with multiple satellite micronuclei (white arrow), indicating a hybrid mitotic catastrophe/mitotic slippage response. **g.** Western blot analyses of the status of Bcl-2 family anti-apoptotic proteins in RR2, Pr1.1, SB2 and GBM6 cells synchronized by double thymidine block and released from the G1-S boundary into vehicle or 1 µM CT-179 for the indicated time points. Actin was included as a loading control. Arrows indicate the time points in CT-179 treated cells when loss of protein is observed. **h-i**. Representative plots for the inhibition of proliferation of primary GBM cell lines after 7 days of treatment with increasing concentrations of CT-179 alone or in combination with S63845 (**h**) or ABT-737 (**i**) as measured by Vialight assay. Data are presented as a percentage of cell viability vs vehicle controls ± s.e.m. Horizontal dashed line = IC50 demarcation, vertical dashed line = 500 nM CT-179 demarcation. **j.** Western blot analyses of cleaved PARP in RR2, Pr.1.1 and GBM6 cells synchronized by double thymidine block and released from the G1-S boundary into vehicle, 1 µM CT-179, 1 µM S63845, 500 nM ABT-737 or combinations of these for 24 h. Actin was included as the loading control.

Staining for the mitotic marker pSer10-Histone H3 (p-H3) likewise showed heterogeneous responses: some cell lines accumulated p-H3 progressively, while others displayed an initial accumulation followed by rapid loss of signal (Fig. 3b). These data indicate that some lines undergo durable mitotic arrest followed by apoptosis, whereas others transiently pause in mitosis and re-enter the cell cycle before eventually becoming apoptotic.

To define the molecular basis of these responses, we synchronized cells at the G1–S boundary by double thymidine block, released them into vehicle or 1 µM CT-179, and monitored Cyclin B1, pThr161-CDK1, and cleaved PARP. Cyclin B1 is induced at mitotic onset, where it associates with CDK1 to drive spindle assembly, and is rapidly degraded after passage through the spindle assembly checkpoint (SAC), with CDK1 inactivated by dephosphorylation^28^. Three distinct response patterns were observed (Fig. 3c). RR2 cells exhibited prolonged accumulation of Cyclin B1 and p-CDK1 from 24 h onward together with cleaved PARP, consistent with mitotic arrest and subsequent apoptosis. SB2 cells displayed normal Cyclin B1 and p-CDK1 turnover at 24 h with no early cleaved PARP, consistent with progression through mitosis without early apoptotic commitment. Pr1.1 cells showed partial reduction of Cyclin B1 and p-CDK1 at 24 h with concomitant cleaved PARP accumulation, indicating a hybrid response in which a subset of cells undergoes mitotic arrest while others complete the cycle. Extended CT-179 exposure (48 h and 72 h) led to loss of total CDK1, CDK2, and Rb protein without changes in their transcripts, again consistent with a translation-level effect (Fig. 2g).

These molecular phenotypes are consistent with mitotic catastrophe in some lines, an intrinsic mechanism in which cells fail to satisfy the SAC, remain blocked in mitosis, and undergo apoptosis^29,30^, and with mitotic slippage in others, a phenomenon in which cells transiently arrest, re-enter the cell cycle, and survive with grossly abnormal nuclear architecture^31^.

Immunofluorescence analysis with α-tubulin staining confirmed these designations: CT-179–treated RR2 cells displayed condensed chromosomes arrested at prometaphase with formed but misaligned spindles, hallmarks of mitotic catastrophe (Fig. 3d); SB2 cells exhibited malformed nuclear architecture with ancillary nuclear lobes, multiple satellite micronuclei, and defective cytokinesis, consistent with mitotic slippage (Fig. 3e); and Pr1.1 cells showed both phenotypes (Fig. 3f), in line with the hybrid cell-cycle profile in Fig. 3c. Together, these data indicate that CT-179 exerts its anti-tumor activity by disrupting mitotic progression, producing either mitotic catastrophe at prometaphase or mitotic slippage, both of which converge on apoptosis with distinct kinetics.

### BH3 mimetics enhance CT-179-induced apoptosis

We next examined whether anti-apoptotic Bcl-2 family proteins limit the efficacy of CT-179. Western blot analyses of synchronized GBM cells released into 1 µM CT-179 showed that one or both of Bcl-2 and Mcl-1 were reduced relative to vehicle controls, while Bcl-xL was not appreciably affected (Fig. 3g). These observations suggested that residual Bcl-2 family activity might attenuate CT-179–induced apoptosis and that BH3 mimetics could enhance its activity. Combination proliferation assays in patient-derived GBM cell lines tested CT-179 with S63845, a selective Mcl-1 inhibitor^32^, or ABT-737, an inhibitor of Bcl-2, Bcl-xL, and Bcl-w but not Mcl-1^33^. Both combinations produced greater-than-additive anti-proliferative activity in several cell lines (Fig. 3h,i and 3i). The pattern of sensitivity was cell-line dependent, for example, SB2 and RN1 showed enhanced CT-179 efficacy with ABT-737 but not S63845, indicating that the dominant pro-survival Bcl-2 family member differs across GBM lines. In synchronized RR2, Pr1.1, and GBM6 cells, combination of either S63845 or ABT-737 with CT-179 augmented cleaved PARP induction relative to single agents (Fig. 3j), consistent with enhanced apoptotic signaling. These findings support a rationale for combining CT-179 with BH3 mimetics in GBM contexts where Bcl-2 or Mcl-1 limits apoptotic commitment.

### CT-179 depletes OLIG2 and induces apoptosis in patient-derived GBM organoids

To corroborate CT-179 target engagement and activity in a patient-derived three-dimensional model that preserves tumor architecture, we generated glioblastoma organoids from freshly resected tumor tissue (line QBC512), which expressed high OLIG2 mRNA, and treated them with CT-179 across a 0.5 to 4 µM range, selected around the 3.77 µM IC50 determined in the matched adherent QBC512 line, with or without a single 2 Gy radiotherapy (RT) fraction (Supplementary Figure S7). CT-179 dose-dependently reduced organoid area and sphericity, a surrogate for structural integrity (P < 0.0001 at 2 and 4 µM versus control), and depleted OLIG2-positive cells from approximately 37% in controls to approximately 2% at 4 µM (P < 0.0001), confirming in-organoid target engagement. At 4 µM, this was accompanied by an increase in cleaved caspase-3-positive cells, consistent with apoptosis. The addition of RT produced a modest further reduction in sphericity at 4 µM (P = 0.0317) but did not significantly augment OLIG2 loss or, on its own, reach significance for apoptosis, indicating that CT-179 was the dominant driver of the organoid response.

### CT-179 inhibits FOXR2-activated neuroblastoma tumors in vivo

CNS neuroblastoma with FOXR2 activation (CNS NB-FOXR2) is an aggressive pediatric cerebral tumor in which OLIG2 is robustly expressed^34^. To assess the activity of CT-179 against a primary OLIG2-positive tumor in vivo, we used an established zebrafish model of CNS NB-FOXR2 in which oncogenic human NRAS^Q61R^ is expressed in OLIG2-positive OPCs^35^.

Embryonic brain transplantation followed by drug treatment^36^ was used to evaluate tumor responses.

Treatment with 1 µM CT-179 for 6 days produced a significant reduction in tumor burden compared with vehicle controls, with efficacy that exceeded that of the MEK inhibitor selumetinib included as a positive control (Fig. 4a,b and 4b). These data indicate that CT-179 inhibits the growth of a primary OLIG2-driven tumor in vivo and support its activity beyond GBM in OLIG2-dependent CNS malignancies.

**Figure 4:**
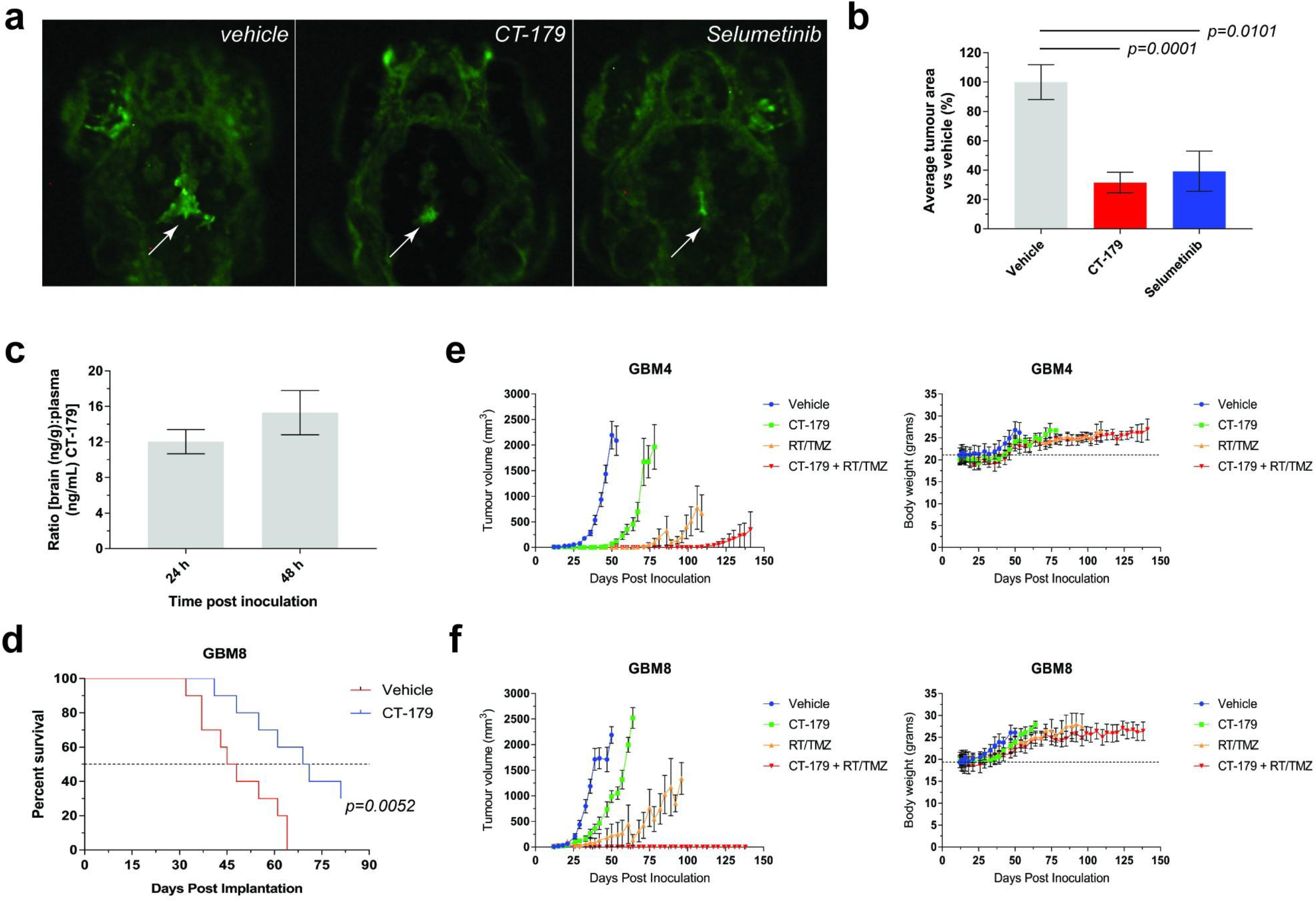
CT-179 crosses the blood-brain barrier, inhibits orthotopic tumor growth and shows enhanced activity when used with existing clinical standard of care. a. Representative whole-mount fluorescent images of day 8 dpf zebrafish avatars expressing GFP-tagged oncogenic N-RAS for visualization in the presence of vehicle, 1 µM CT-179 or 50 µM selumetinib as a positive control. White arrows show GFP-tagged tumors in the 4^th^ ventricle. Background is due to auto-fluorescence of iridophores in the eye. **b.** Quantitation of inhibition of tumor growth in zebrafish avatars after treatment with vehicle, CT-179 or selumetinib. Data are presented as the median % tumor area vs vehicle controls ± s.e.m. Significance values were obtained by one-way ANOVA. **c.** Measurement of the brain to plasma ratios of CT-179 present in the brain parenchyma at 24 h and 48 h after the third of three consecutive once-daily oral gavage doses of 5 mg/kg CT-179. **d.** Kaplan-Meier survival curves demonstrating CT-179 single agent anti-tumor activity against GBM8 orthotopic xenografts in NSG mice when administered by intraperitoneal injection for 14 days qd at 20 mg/kg. Data are presented as the percentage survival of mice in each group over time. A significant 24-day increase in median survival was observed (vehicle: 46 days vs CT-179: 70 days) after log-rank (Mantel-Cox) analysis. Single-agent CT-179 activity in adult MGMT-unmethylated G06 orthotopic xenografts (Kaplan-Meier curve; 150 mg/kg/day oral; ≈46% median survival extension) is shown in Supplementary Figure S2. **e-f.** Tumor growth plots (left graphs) and body weight plots (right graphs) for female NSG (NOD.Cg-Prkdc^scid^ Il2rg^tm1Wjl^/SzJ) mice (The Jackson Laboratory) implanted subcutaneously with GBM4 (**e**) or GBM8 (**f**) cell lines and subsequently treated with vehicle, CT-179, RT/TMZ or a combination of both. The combination of CT-179 with RT/TMZ in the GBM8 xenograft model completely eliminated all the xenografts and was significantly more efficacious than all other groups. In the GBM4 model, the CT-179 with RT/TMZ combination produced the greatest tumor reduction, completely eliminating 7 out of 8 xenografts, although the difference from RT/TMZ alone did not reach statistical significance (P = 0.052; ‡). Data are presented as tumor volume (mm^3^) over time (left graphs) or body weight in grams over time (right graphs). * = comparison of CT-179 alone, RT/TMZ alone and combination to vehicle by one way ANOVA at Day 50 for GBM4 and Day 42 for GBM8, *P* < 0.001 for both tests; † = comparison of RT/TMZ and combination with CT-179 alone by one way ANOVA at Day 74 for GBM4 and Day 61 for GBM8, *P* < 0.001 for both tests; ‡ = comparison of combination to RT/TMZ alone by student’s t-test at Day 109 for GBM4 (*P* = 0.052) and Day 85 for GBM8 (*P* = 0.0076).

### CT-179 crosses the blood-brain barrier to potently inhibit GBM tumor growth

We next assessed whether CT-179 reaches therapeutic concentrations in the brain. A single 20 mg/kg oral gavage dose in C57BL/6J mice indicated an oral bioavailability of approximately 76% and a plasma half-life of 10.7 h that on its own would not justify once-daily dosing. Daily oral dosing at 5 mg/kg produced sustained CNS exposure with brain-to-plasma ratios (brain concentrations in ng/g taken as equivalent to plasma concentrations in ng/mL) of approximately 12 at 24 h and 15 at 48 h after the third dose (Fig. 4c), consistent with efficient blood-brain barrier penetration and sustained brain retention that supports once-daily oral dosing. Intraperitoneal administration produced a similar brain-exposure profile, with brain-to-plasma ratios rising over 24 h to approximately 10–15, indicating that CNS exposure of CT-179 is comparable following oral and intraperitoneal dosing (Supplementary Figure S6).

In an orthotopic GBM8 patient-derived xenograft model in NSG mice, CT-179 administered by intraperitoneal injection at 20 mg/kg once daily for 14 days as a single agent produced a significant delay in tumor progression and extended median survival by 24 days relative to vehicle controls (70 vs. 46 days; *P* = 0.0052; Fig. 4d). Immunohistochemistry of resected tumor tissue from these animals showed reductions in nuclear OLIG2 and Ki-67 staining (Supplementary Figure S5), consistent with in-tumor target engagement and reduced proliferation. To characterize the oral dose-response, we administered CT-179 by oral gavage at 1 or 5 mg/kg once daily to NSG mice bearing orthotopic GBM8 xenografts. Daily oral CT-179 at 5 mg/kg significantly extended median survival (approximately 42% benefit relative to vehicle; Supplementary Figure S4), whereas the 1 mg/kg dose was not significant. Daily 150 mg/kg oral CT-179 also significantly extended median survival in adult MGMT-unmethylated G06 intracranial xenografts (approximately 46% benefit; Supplementary Figure S2), establishing single-agent activity across two independent adult orthotopic backgrounds.

### CT-179 potently enhances standard-of-care therapy

Because CT-179 induces profound mitotic and chromosomal instability, we hypothesized that it would enhance the efficacy of GBM standard of care, namely radiotherapy combined with temozolomide (RT/TMZ). We tested CT-179 alone, RT/TMZ alone, and the combination in subcutaneous GBM4 and GBM8 patient-derived xenografts in NSG mice; both lines harbor *MGMT* promoter methylation (95% and 55% by pyrosequencing, respectively), a context in which TMZ is expected to contribute meaningfully to standard-of-care activity.

Single-agent CT-179 and RT/TMZ each delayed tumor growth in both models (Fig. 4e and 4f, left panels). The combination of CT-179 with RT/TMZ produced markedly greater tumor control than either treatment alone: in the GBM8 model, the combination eliminated all xenografts and was significantly more efficacious than every other arm, including RT/TMZ alone (P = 0.0076 at Day 85); in the GBM4 model, the combination eliminated 7 of 8 xenografts, although the combination-versus-RT/TMZ contrast did not reach statistical significance (P = 0.052 at Day 109). The GBM4 combination thus represents a strong trend that reached statistical significance in GBM8. Body weights remained comparable to vehicle controls across all treatment groups in both studies (Fig. 4e and 4f, right panels), indicating that the combination was well tolerated.

Together, these data demonstrate that CT-179 crosses the blood-brain barrier at concentrations sufficient for single-agent activity and substantially enhances the efficacy of standard-of-care chemoradiotherapy in patient-derived GBM models.

### CT-179 enhances radiotherapy in an orthotopic MGMT-unmethylated model

To determine whether the chemoradiosensitization observed in flank xenografts extends to intracranial disease in a setting where temozolomide contributes minimally, we tested CT-179 in combination with radiotherapy in an MGMT-unmethylated orthotopic patient-derived model. In intracranial GBM6 xenografts, an MGMT-unmethylated line in which CT-179 monotherapy did not extend median survival relative to vehicle (35 vs. 32 days; *P* = 0.21), addition of CT-179 (50 mg/kg intraperitoneally, three times weekly for three weeks) to fractionated radiotherapy (2 Gy × 4 fractions over 8 days; 8 Gy total) significantly extended median survival relative to RT alone (48 vs. 38.5 days; *P* = 0.0248; Fig. 5). These data indicate that CT-179 enhances radiotherapy activity in an orthotopic, MGMT-unmethylated patient-derived xenograft, the population for which temozolomide contributes least.

**Figure 5:**
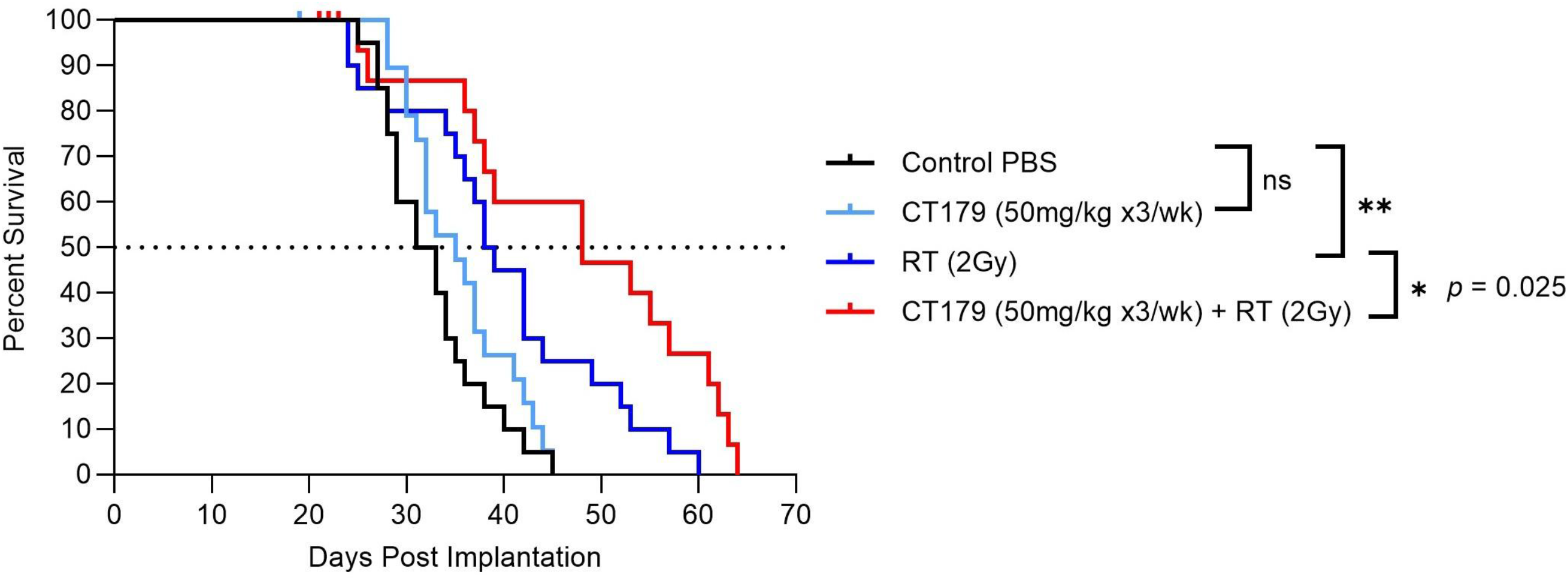
CT-179 enhances radiotherapy in MGMT-unmethylated orthotopic GBM. Kaplan-Meier survival curves for NOD-RAG mice bearing intracranial GBM6 (MGMT-unmethylated) patient-derived xenografts (1 × 10⁶ cells per implant) randomized (vehicle, n = 20; CT-179, n = 19; RT, n = 20; CT-179 + RT, n = 15) to vehicle, CT-179 alone (50 mg/kg intraperitoneally, three times weekly for three weeks beginning Day 8), fractionated radiotherapy (2 Gy × 4 fractions over 8 days; 8 Gy total), or the combination of CT-179 + RT. CT-179 monotherapy did not extend median survival (CT-179: 35 days vs vehicle: 32 days; *P* = 0.21; not significant). Combination of CT-179 with RT significantly extended median survival relative to RT alone (CT-179 + RT: 48 days vs RT: 38.5 days; *P* = 0.0248 by log-rank test). Body weights remained comparable to vehicle controls across treatment groups.

## Discussion

GBM remains uniformly fatal under current standard of care, and disease-modifying therapies that disable the lineage transcriptional programs sustaining the GBM stem-like state have remained elusive. The data presented here address this knowledge gap by characterizing CT-179, a brain-penetrant small molecule that engages OLIG2 within tumor cells, in adult GBM. We show that CT-179 binds OLIG2 in patient-derived GBM cells, alters its conformation on chromatin without preventing dimerization or fully displacing it from DNA, depletes OLIG2-positive OPCs in vivo, induces apoptosis across a phenotypically diverse panel of patient-derived GBM lines, achieves brain exposures sufficient for single-agent activity in orthotopic xenografts, and substantially enhances the efficacy of RT/TMZ. Patient-derived GBM organoids corroborated these findings, with CT-179 dose-dependently depleting OLIG2 and inducing apoptosis in a three-dimensional model that preserves tumor architecture (Supplementary Figure S7). Together, these findings establish proof of concept for direct pharmacologic inhibition of OLIG2 as a therapeutic strategy in GBM.

The mode of OLIG2 engagement by CT-179 is mechanistically distinct from conventional approaches to transcription factor inhibition. CETSA in patient-derived GBM cells and differential scanning fluorimetry on a recombinant bHLH construct converge on the same conclusion: CT-179 binds an OLIG2•dsDNA complex and produces a small but reproducible destabilization (1.8°C) of the dimer-DNA species without displacing OLIG2 from chromatin or preventing dimer formation, as confirmed by DSS crosslinking and EMSA. This conformational mode of action is consistent with the in silico pharmacophore strategy that originally targeted the OLIG2 dimerization interface as a tractable surface^12,13^ and complements, rather than recapitulates, prior genetic and post-translational studies in which OLIG2 phosphorylation status and lineage-restricted transcriptional output govern its proliferative and tumor-promoting activity^6,7^. A recent study using fluorescence cross-correlation spectroscopy (FCCS) on full-length OLIG2 in live HEK-293 cells reported that CT-179 impedes OLIG2 dimerization without reducing OLIG2 protein abundance^37^. We do not regard this as inconsistent with the present data: FCCS measures the kinetic equilibrium of full-length OLIG2 dimerization in the cellular environment and is sensitive to transient or partial dissociation that DSS crosslinking and end-point EMSA, both of which capture stably formed complexes, may not detect. Together, these observations indicate that CT-179 likely destabilizes the OLIG2 dimer-DNA species in a manner detectable by sensitive live-cell biophysics while leaving a substantial fraction of crosslink-stable, DNA-bound dimer intact in standard biochemical assays.

Functional consequences of this engagement converge on disrupted mitotic progression. We observed three distinct cell-line responses to CT-179: durable mitotic arrest with prolonged Cyclin B1 and pThr161-CDK1 accumulation followed by apoptosis^29,30^, transient arrest followed by re-entry into the cell cycle with grossly abnormal nuclear architecture^31^, and a hybrid pattern that combines both. Importantly, sustained activation of ERK1/2, AKT, and STAT3 in CT-179-treated cells indicates that the anti-tumor effect is not explained by collapse of canonical RTK-driven survival signaling, despite a translation-level reduction in EGFR, PDGFR-α, and HER4 protein. The reduction in RTK protein, which occurs without a corresponding decrease in transcript abundance and parallels a reduction in global nascent translation, is best interpreted as a downstream consequence of OLIG2 inhibition rather than its proximate driver of cell death, although this ordering is inferred from the concurrent persistence of ERK1/2, AKT, and STAT3 signaling rather than directly demonstrated. This interpretation is consistent with prior reports that OLIG2 enforces a lineage-restricted RTK signature in glioma stem-like cells^9,10^ but extends those observations by showing that a chemically tractable OLIG2 inhibitor uncouples RTK protein abundance from downstream pathway activation while still producing cytotoxicity.

The therapeutic implications of these mechanisms inform combination strategy. Because cells executing mitotic catastrophe or slippage remain dependent on residual anti-apoptotic Bcl-2 family activity to delay apoptotic commitment, we asked whether BH3 mimetics would lower the threshold for cell death. CT-179 reduced Bcl-2 and/or Mcl-1 protein in synchronized GBM cells, and combination with the Mcl-1 selective inhibitor S63845^32^ or the Bcl-2/Bcl-xL/Bcl-w inhibitor ABT-737^33^ produced greater-than-additive anti-proliferative activity in a cell-line-dependent manner. The dependence on a specific Bcl-2 family member (for example, ABT-737 sensitivity in SB2 and RN1) suggests that a Bcl-2 family expression readout could serve as a candidate biomarker for selecting CT-179 combination partners. Independently, the combination of CT-179 with RT/TMZ eliminated all GBM8 xenografts and 7 of 8 GBM4 xenografts without weight loss, providing a rationale to pair CT-179 with the existing GBM standard of care rather than to position it solely as a monotherapy. We note that brain accumulation of CT-179 after oral dosing may be context-dependent: in MYC-amplified medulloblastoma orthotopic xenografts, oral CT-179 was reported to be insufficient to suppress intracranial tumor growth, and intrathecal delivery was required to achieve durable combination activity with craniospinal irradiation^38^. In our studies, oral CT-179 produced steady accumulation in brain parenchyma in C57BL/6J mice in pharmacokinetic experiments, while intraperitoneal CT-179 at 20 mg/kg/day for 14 days conferred significant survival benefit in the orthotopic GBM8 model and oral CT-179 at 70 mg/kg/day combined with RT/TMZ produced near-complete tumor clearance in subcutaneous GBM4 and GBM8 models. A recent independent study in pediatric high-grade glioma similarly reported that oral CT-179 at 24 mg/kg/day for 5 days achieves measurable accumulation across cerebrum, cerebellum, brain stem, and patient-derived orthotopic xenograft tumors, and at 200 mg/kg every 4–5 days potentiates fractionated radiotherapy in 2 of 4 pediatric GBM models^39^, supporting brain penetration of orally administered CT-179 in adult and pediatric glioma. These observations indicate that the route and dose of administration required for adequate tumor exposure will need to be optimized for each indication and tumor location. This combination rationale is reinforced by the observation that OLIG2 opposes p53-dependent responses to genotoxic stress^7^: pharmacologic inhibition of OLIG2 should, in principle, restore the genotoxic response to RT/TMZ in OLIG2-positive tumors.

The orthotopic combination data carries a translational implication of particular interest in MGMT-unmethylated disease. In intracranial GBM6 xenografts, CT-179 monotherapy did not change median survival relative to vehicle, but combination with radiotherapy produced a significant survival benefit over radiotherapy alone. The contrast between the two MGMT-unmethylated orthotopic models is instructive: single-agent CT-179 extended survival in G06 (150 mg/kg/day orally) but not in GBM6 (50 mg/kg intraperitoneally, three times weekly), indicating that the divergent monotherapy outcomes more likely reflect differences in dose and achieved tumor exposure than MGMT methylation status. Notably, GBM6 is among the in-vitro CT-179-sensitive lines (IC50 < 360 nM; Fig. 2a), so its lack of single-agent activity in the orthotopic setting is most consistent with insufficient tumor exposure at the dose and schedule used rather than intrinsic resistance. Because MGMT methylation is the principal determinant of temozolomide benefit and OLIG2 opposes p53-dependent genotoxic responses, pharmacologic OLIG2 inhibition may sensitize MGMT-unmethylated tumors to radiotherapy via restored genotoxic-stress signaling. This motivates an MGMT-stratified clinical hypothesis pairing CT-179 with radiotherapy in MGMT-unmethylated GBM, the population for which temozolomide contributes least and consistent with the OPAL Treatment Arm 2 design^16,39,40,41^. Because oral and intraperitoneal dosing produced comparable brain exposure (Supplementary Figure S6), the combination benefit observed with intraperitoneal CT-179 in GBM6 would be expected to be reproducible with oral administration at an equivalent exposure.

Recent independent work also illuminates additional combination strategies and the cellular subpopulations that determine CT-179 sensitivity. Single-cell transcriptomic analyses of GBM tumors integrated with drug response signatures predict that CT-179 preferentially targets oligodendrocyte-progenitor-like (OPC-like) and neural-progenitor-like (NPC-like) tumor cell states while leaving astrocyte-like (AC-like) cells relatively resistant^42,43^; pairing CT-179 with the anti-EGFR antibody-drug conjugate Depatuxizumab Mafodotin (Depatux-M, ABT-414), which targets EGFR-expressing AC-like cells, significantly extends survival in orthotopic GBM patient-derived xenografts^44^. This single-cell-resolution view of complementary targeting is consistent with our observation that CT-179 reduces stem-like populations and depletes specific RTK protein pools without abolishing downstream pathway activation, and provides an additional combination rationale beyond standard-of-care chemoradiotherapy. CT-179 has also recently been shown to disrupt the immunosuppressive tumor microenvironment of GBM: OLIG2 represses CXCL10 transcription through HDAC7 recruitment, suppressing CD8+ T cell infiltration and activation, and pharmacologic inhibition of OLIG2 with CT-179 sensitizes GBM tumors to anti-PD-L1 therapy in immunocompetent models^11,40^. Together, these findings suggest that CT-179 may have value not only as a chemoradiosensitizer but also as a means of converting immunologically “cold” GBM tumors into a state that is more responsive to immune checkpoint blockade.

Considered alongside independent reports, CT-179 has now been shown to confer in vivo survival or tumor-burden benefit across a structurally diverse set of OLIG2-dependent CNS malignancies, including adult GBM (this study), pediatric high-grade glioma in combination with fractionated radiotherapy, SHH-subgroup medulloblastoma^16,37^, MYC-amplified medulloblastoma^38^, and zebrafish CNS NB-FOXR2 orthotopic xenografts^34,35^. The convergence of in vivo activity across this range of OLIG2-driven tumor backgrounds, in independent laboratories using independent models, indicates that the therapeutic relevance of OLIG2 inhibition extends beyond adult GBM and supports OLIG2 as a tractable, broadly relevant target in CNS oncology. The same logic is supported by independent reports that OLIG2 is required for proliferation of patient-derived diffuse intrinsic pontine glioma (DIPG; reclassified under the 2021 WHO classification of CNS tumors as H3 K27-altered diffuse midline glioma, DMG^3^) lines and is associated with significantly shorter overall survival in OLIG2-positive DIPG^45^, and by evidence that OLIG2 confers radiation and chemoresistance and orchestrates a proneural transcriptional program whose disruption shifts tumors toward a mesenchymal phenotype^46^. The selective reduction of OPCs in zebrafish embryos without overt disruption of mature oligodendrocytes is consistent with the lineage-restricted CNS expression of OLIG2 in adults^5^ and supports a tractable therapeutic window, although chronic CNS exposure in mature animals will require dedicated assessment. Several limitations should be acknowledged. The data are entirely preclinical, and clinical activity in patients with GBM remains to be established; pharmacodynamic biomarkers of OLIG2 engagement in tumor tissue will be required to interpret early-phase trials. The patient-derived cell line panel, although diverse, does not span every molecular subtype of GBM, and predictors of CT-179 sensitivity (including the determinants of mitotic catastrophe versus slippage and the cell-line-specific Bcl-2 family dependency) warrant further investigation. Finally, the durability of orthotopic responses and the potential for resistance after extended exposure remain open questions.

Taken together, the data position CT-179 as a brain-penetrant, mechanistically novel inhibitor of an established GBM driver, with single-agent activity in orthotopic models and substantial enhancement of standard-of-care chemoradiotherapy. These findings support the further development of CT-179 for GBM and motivate translational studies in other OLIG2-dependent CNS malignancies.

## Conclusions

CT-179 is a brain-penetrant small-molecule inhibitor of OLIG2 that engages its target on chromatin, alters dimer-DNA conformation without displacing OLIG2 from its binding sites, and converts the OLIG2-driven proliferative program of glioblastoma into apoptotic cell death through mitotic catastrophe and slippage. Oral CT-179 extends single-agent survival in two independent adult orthotopic patient-derived backgrounds, and intraperitoneal CT-179 is associated with reduced intratumoral OLIG2 and Ki-67 staining in GBM8 tumors; combination with radiotherapy and temozolomide eliminates the great majority of subcutaneous xenografts without overt toxicity, and significantly extends survival when combined with radiotherapy in MGMT-unmethylated GBM6 orthotopic xenografts where temozolomide contributes minimally. Considered alongside independent reports of in vivo activity in pediatric high-grade glioma, SHH- and MYC-amplified medulloblastoma, and CNS NB-FOXR2 xenografts, the cumulative survival evidence across structurally diverse OLIG2-dependent CNS malignancies provides robust target validation for OLIG2 in CNS oncology. These findings establish proof of concept for direct pharmacologic inhibition of a previously undruggable transcription factor in glioblastoma, identify rational combination partners including BH3 mimetics and standard-of-care chemoradiotherapy, and motivate an MGMT-stratified clinical strategy in which CT-179 is paired with radiotherapy in MGMT-unmethylated GBM. The principal next step is clinical evaluation of CT-179 in glioblastoma, supported by pharmacodynamic biomarkers of OLIG2 engagement and by parallel translational assessment in other OLIG2-dependent central nervous system malignancies.

## Funding

T.G.J. is a Principal Research Fellow with the National Health and Medical Research Council of Australia. D.J.G. is supported by grants from the National Health and Medical Research Council of Australia (1103068), the U.S. Department of Defense (CA150132), and Cancer Council Victoria (1145028). G.D.S., G.A., S.K. (Knowles), and G.B. research was funded in part by the Cancer Prevention and Research Institute of Texas (CPRIT) contract award DP140034. R.A.S. was supported by the Huntsman Cancer Foundation and the National Institutes of Health/National Cancer Institute (P30CA042014). R.M. acknowledges financial support from the American Brain Tumor Association. The in vivo pharmacology and pharmacokinetic studies of CT-179 and the associated medicinal-chemistry program were sponsored by Curtana Pharmaceuticals, Inc.

## Acknowledgements

We thank Caroline Humphries, Rahul Kollipara and the McDermott Next-Generation Sequencing Core (UT Southwestern Medical Center) for assistance with the RNA-seq studies. During the preparation of this manuscript, the authors used a generative artificial-intelligence tool (Claude Opus 4.8) to assist with drafting and language editing. All AI-assisted text was reviewed, verified, and edited by the authors, who take full responsibility for the integrity, accuracy, and final content of the work. No AI tool is listed as an author.

## Author Contributions

Author contributions are described per the CRediT taxonomy. Conceptualization: T.G.J., S.K. (Kesari), G.D.S., G.A., G.B. Methodology: S.A.G., S.C.C., G.A., R.B., M.C., I.B., Z.C.B., D.J.G. (Garama), R.M., D.J.G. (Gough). Investigation: S.A.G., S.C.C., R.B., M.C., I.B., Z.C.B., D.J.G. (Garama), R.M., A.K., R.K., B.S. Resources / Medicinal Chemistry: G.B., F.C.T., S.B.R., A.L., S.K. (Knowles). Formal Analysis: S.A.G., S.C.C., G.A., R.A.S., D.J.G. (Gough), A.K., R.K. Data Curation: A.K., R.K. Visualization: S.A.G., S.C.C., G.A. Writing – Original Draft: S.A.G., S.C.C., G.A., G.B., G.D.S., R.A.S., T.G.J. Writing – Review & Editing: all authors. Supervision: T.G.J., S.K. (Kesari), R.A.S., B.W.D., D.J.G. (Gough), G.D.S. Funding Acquisition: T.G.J., D.J.G. (Garama), G.D.S., R.A.S., R.M. Project Administration: T.G.J. S.A.G. and S.C.C. contributed equally to this work.

## Conflict of Interest

During the conduct of this work, GDS, GB, GA, and SK (Knowles) were employees of Curtana Pharmaceuticals, Inc.; GDS also served as an executive officer and a member of the Board of Directors. SK (Kesari) is a co-founder, scientific advisor, and member of the Board of Directors of Curtana. FCT and SBR are co-founders and/or scientific advisors of Curtana. AL performed medicinal-chemistry work as a graduate student at the Center for Innovative Drug Discovery, The University of Texas at San Antonio, which acted as a contract research provider to Curtana. TJ and BD were academic investigators with no financial interest in Curtana at the time the work was performed; both were subsequently granted stock options. GDS, SK (Knowles), FCT, SBR, GB, GA, AL, TJ, BD, and SK (Kesari) hold equity (shares and/or stock options) in Curtana Pharmaceuticals, Inc. CT-179 is proprietary to Curtana Pharmaceuticals, Inc., which sponsored the in vivo studies described here. The remaining authors declare no competing interests.

## Data Availability Statement

The RNA-sequencing data generated in this study (36 libraries from the C419, CD133, and GBM4 models) have been deposited in the NCBI Gene Expression Omnibus (GEO) and are publicly available under accession number GSE336473 as of the date of publication. All other data supporting the findings of this study are available within the article and its supplementary information. Additional information and unprocessed data files are available from the corresponding author upon reasonable request. CT-179 was developed by Curtana Pharmaceuticals, Inc.; requests for the compound should be directed to Curtana Pharmaceuticals.

## Online Methods

### Animal ethics and reporting

All vertebrate animal procedures were performed at institutions or contract research organizations operating under formally approved animal care and use programs and were conducted in accordance with the relevant national and institutional guidelines in effect at the time. The orthotopic GBM8 xenograft study was performed at the University of California, San Diego, under approval from the UCSD Institutional Animal Care and Use Committee (IACUC). The subcutaneous GBM4 and GBM8 flank xenograft studies were performed by Charles River Discovery Services (Morrisville, NC, USA), an AAALAC-accredited contract research organization, on behalf of Curtana Pharmaceuticals, Inc., under Charles River Discovery Services’ institutional animal care and use program. Pharmacokinetic and brain-accumulation studies in C57BL/6J mice were performed at BioDuro (Beijing, China) on behalf of Curtana Pharmaceuticals, Inc., under BioDuro’s institutional animal care and use program; the intraperitoneal brain-accumulation study was conducted under IACUC protocol BBD-201503014. Zebrafish studies were performed at the Huntsman Cancer Institute (University of Utah) under approval from the University of Utah Institutional Animal Care and Use Committee (IACUC #16-03019). With the exception of the Utah zebrafish program and the BioDuro intraperitoneal brain-accumulation study noted above, the specific institutional protocol numbers issued at the time of the studies are no longer accessible from contemporaneous records, given the period that has elapsed since the work was conducted; the conducting institutions and the relevant approval frameworks are named here in lieu of those numbers. Reporting in this manuscript follows the ARRIVE 2.0 guidelines to the extent permitted by the historical nature of several studies, and a completed ARRIVE 2.0 checklist accompanies this submission with non-retrievable items marked accordingly. As detailed below, certain items could not be reconstructed from contemporaneous records, including certain institutional protocol numbers, formal sample-size justification, and formally documented randomization and blinding procedures. Animals were group-housed under standard 12 h light/12 h dark cycles with ad libitum access to food and water. Sample size for in vivo efficacy studies (n = 8 per group for subcutaneous xenograft studies; for orthotopic survival studies, n = 10 for the GBM8 single-agent study, n = 7–8 for the GBM8 oral dose-response study, n = 7 for the G06 study, and n = 15–20 across arms for the GBM6 combination study) was selected based on prior experience with comparable patient-derived xenograft models in our laboratories and was not formally derived from a power calculation. Animals were randomized to treatment groups by tumor volume immediately prior to first dose; tumor volumes were measured by an investigator blinded to treatment assignment. Inclusion criteria required successful tumor engraftment by day 12 (subcutaneous models) or by day 7 (orthotopic models); animals failing engraftment, presenting with poor body condition unrelated to tumor burden, or showing technical complications during inoculation were excluded prior to randomization. No animals were excluded after randomization. Specific randomization and blinding methods were performed at the time of the studies but were not formally documented in standard operating procedures available at the time; the descriptions provided here reflect the laboratory practices in effect during the conduct of the work.

### Human tissue collection and ethics

Tumor tissue was collected from patients undergoing surgery at the Royal Brisbane and Women’s Hospital (RBWH) or the Queensland Children’s Hospital with informed consent and human ethics approval from the QIMR Berghofer Medical Research Institute (P3420) and RBWH human research ethics committees (HREC/17/QRBW/577).

### Reagents

ViaLight™ Plus cell proliferation and cytotoxicity bioassay kits were from Lonza (Basel, Switzerland). Caspase-Glo® 3/7 assays were from Promega (Madison, WI, USA). RNeasy® mini kit (250) was from QIAGEN (Hilden, Germany). Protein A/G agarose beads were from Santa Cruz Biotechnology (Dallas, TX, USA). Odyssey® infrared EMSA kits were from LI-COR (Lincoln, NE, USA). The following reagents were all from Thermo Fisher Scientific (Waltham, MA, USA): FxCycle™ PI/RNase staining solution, Pierce® RIPA lysis buffer, Halt® protease and phosphatase inhibitor cocktail, BCA assay, NuPAGE™ 4-12% Bis-Tris protein gel, iBlot™ transfer stack-PVDF (mini or regular), NE-PER™ nuclear and cytoplasmic extraction reagents kit, 16% paraformaldehyde (PFA), TaqMan® gene expression assays for *Olig2, Cdk1, Cdk2, Cdk6, Cdk7, Cdk4, Rb, Pdgfra, Met, Egfr, Erbb4* and *Actb*, SlowFade™ diamond antifade mountant with DAPI and SuperScript® III first-strand synthesis supermix kit.

### Antibodies

OLIG2 rabbit antibody (#ab136253) was from Abcam (Cambridge, UK). OLIG2 mouse mAb, clone 211F1.1 (#MABN50), CDK6 mouse antibody (#MABC280) were from Merck (Kenilworth, NJ, USA). Actin mouse mAb (#ACTN05 (C4)), secondary goat-anti rabbit (AF800 conjugate), secondary goat-anti mouse (AF680 conjugate), alpha-tubulin mAb (AF488 conjugate) (#322588), Annexin V (AF647 conjugate) (#A23204) were from Thermo Fisher Scientific (Waltham, MA, USA). CDK2 mouse mAb (#sc6248) and ERBB4 rabbit antibody (#sc283) were from Santa Cruz Biotechnology. mAb806 (EGFR antibody) was developed and described previously^47^. Herceptin was a kind gift from Dr. Tim Adams, CSIRO, Melbourne, Australia. The following antibodies were all from Cell Signaling Technology (Danvers, MA, USA): PDGFR-α rabbit mAb (#3174), p-AKT (Ser473) rabbit mAb (#4060), AKT mouse mAb (#2920), p-ERK1/2 (Thr202/Tyr204) rabbit mAb (#4370), ERK1/2 mouse mAb (#4696), p-STAT3 (Tyr705) mouse mAb (#9138), STAT3 rabbit mAb (#4904), Cyclin B1 rabbit mAb (#12231), p-CDK1 (Thr161) antibody (#9114), CDK1 mouse mAb (#9116), Mcl-1 rabbit mAb (#94296), MET mouse mAb (#3148), p-CDK2 (Thr160) antibody (#2561), p-RB (Ser807/811) rabbit mAb (#8516), RB mouse mAb (#9309), p-CDK6 (Tyr24) rabbit antibody (#sc293097), p-histone H3 (Ser10) rabbit mAb (AF647 conjugate) (#3377), cleaved-PARP rabbit mAb (#5625), Bcl-2 (124) mouse mAb (#15071) and Bcl-xL (54H6) rabbit mAb (#2764).

### Synthesis of CT-179

The synthesis of CT-179 is described in detail in our issued patent US 11,691,951 B2^15^.

### Primary GBM serum-free cell culture

RR2, Pr1.1, SB2, RN1, WK1, GBM6, GBM39, BAH1, GBML1, and GBML2 were obtained by our laboratory as previously described^23,24^. GBM9 and SK748 were kindly donated by Dr. Amyn Habib (University of Texas Southwestern Medical Center, Dallas, TX, USA). Of these, the eleven lines screened for CT-179 anti-proliferative activity in Fig. 2a were RR2, Pr1.1, SB2, RN1, WK1, GBM6, GBM39, GBML1, GBML2, GBM9, and SK748. All of the primary patient-derived GBM cell lines were grown as spheres or on Matrigel coated plates as previously described^23^. The QBC512 line used for the organoid studies was established from Cavitron Ultrasonic Surgical Aspirator (CUSA) aspirate-derived tissue fragments as reported previously23: fragments were pelleted, red blood cells were removed by incubation in RBC lysis buffer, and tissue was washed in PBS, mechanically and enzymatically dissociated, and seeded for culture. QBC512 was maintained as a glioma neural stem (GNS) cell culture in DMEM/F12 knockout medium supplemented with 1× StemPro Neural Supplement (Gibco), 1× GlutaMAX Supplement (Gibco), 20 ng/mL EGF, and 20 ng/mL FGFb, with 100 U/mL penicillin and 100 µg/mL streptomycin, and grown adherently on basement-membrane Matrigel matrix. The medulloblastoma line DAOY, used as an OLIG2-positive control, was obtained from the American Type Culture Collection (ATCC) and cultured in RPMI supplemented with 1% penicillin-streptomycin, 1% GlutaMAX-I CTS (Gibco), and 10% fetal bovine serum (Gibco). All experiments used cell lines between passages 1 and 25.

### Low-adhesion culture plates

Cell culture plates (Corning) were rinsed with an anti-adherence rinsing solution (STEMCELL Technologies) for 30 minutes and washed with sterile PBS. PBS was aspirated and plates were used immediately for organoid culture.

### Glioma organoid generation, culture, and biobanking

Glioma organoids were generated, cultured, and maintained according to Jacob et al. [REF-JACOB — insert Mendeley citation]. Briefly, a small piece of tumor tissue (approximately 100 mg) was taken from the periphery of the resected sample, avoiding necrotic areas, and immediately transferred into cold glioblastoma dissection medium containing Hibernate A before micro-dissection. Tumor samples were processed within approximately 6 hours of surgical resection, including cutting into small pieces with surgical scissors, followed by washing with PBS and DMEM/F12 medium. Twenty to sixty tumor pieces per well were cultured in 6-well ultra-low-attachment plates in GBO medium on a CO₂-resistant orbital shaker rotating at 120 rpm within a sterile incubator at 37 °C and 5% CO₂.

The day of tumor dissection was designated day 0. Successful organoid establishment was defined by rounding of micro-dissected fragments within the first week (±1 day), with non-viable or poor-quality pieces removed. Viability was assessed by rounding and bright transparency under phase-contrast microscopy. For experimental assays, organoids were not passaged by bisection more than twice and were no older than 2 weeks at the start of an assay, to maintain consistency and avoid changes in growth characteristics.

### Irradiation of organoids

Organoids were transferred into 24-well plates, treated with CT-179, and, after 90 minutes of incubation, irradiated using a 137Cs gamma-ray source to a dose of 2 Gy (MDS Nordion Gammacell Irradiator).

### Proliferation assays

Proliferation assays were conducted using the ViaLight™ Plus Cell Proliferation and Cytotoxicity BioAssay Kit (Lonza) as previously described^23^ using the indicated concentrations of drugs.

For the adherent QBC512 dose-response assay, cells were seeded in two dimensions in 96-well Matrigel-coated plates, grown to approximately 80% confluency overnight, and treated with CT-179 for 72 h. Medium-only negative controls were included. Cells were fixed in methylated spirits and viability was measured by sulforhodamine B (SRB) assay. Absorbance was measured at 564 nm, and values normalized to the mean of control wells were used to calculate percent cytotoxicity.

### Flow cytometry

Cells were treated with vehicle or 1 µM of CT-179 for the indicated times before dissociation with Accutase and a single wash with ice-cold PBS. For p-Ser10 Histone H3 staining, cells were fixed in 4% PFA for 15 min followed by quenching in TBS for 2 min before permeabilization and blocking in NSC SFM medium containing 0.1% Tween-20 and 0.3% Triton X-100 (NSC SFM TTX) for 1 h. Cells were then stained using AF647-conjugated p-histone H3 rabbit mAb (Cell Signaling) diluted 1:50 in NSC SFM TTX for 3 h at 22°C before washing and final resuspension in NSC SFM TTX. For Annexin V staining, cells were stained using AF647-conjugated Annexin V and Annexin V binding buffer (Life Technologies) according to the manufacturer’s instructions. All tests were run on a FACSCanto II (BD) machine and the data were analyzed using FlowJo^®^ software.

### Cell cycle analysis

Cells were treated with vehicle or 1 μM of CT-179 for 72 h before dissociation with Accutase and fixation in 100% ice-cold ethanol while vortexing. DNA was stained for 15 min at 22°C using FxCycle™ PI/RNase Staining Solution (Thermo Scientific) and analyzed on FACSCanto II (BD). The data were analyzed using FlowJo^®^ software.

### Immunoprecipitation, SDS-PAGE and Western blotting

Cells were lysed on ice for 10 min in RIPA lysis buffer (Thermo Scientific) supplemented with 2 × Halt® protease and phosphatase inhibitor cocktail and 1 mM EDTA (Thermo Scientific). The cells were then centrifuged at 14,000 rpm at 4°C for 20 min, and lysates were collected. After protein concentration determination using the BCA assay (Thermo Scientific), equal amounts of protein were mixed with reducing LDS sample buffer, boiled at 95°C for 5 min, loaded onto 4-12% Bis-Tris SDS-PAGE gels (Thermo Scientific) and run at 150 V. Proteins were transferred to PVDF membranes using an iBlot™ apparatus (P3 for 7 min) and then blocked in Odyssey® blocking buffer (LI-COR) for 1 h. Membranes were probed for target using relevant primary antibodies diluted in Odyssey buffer overnight at 4°C with rocking. Blots were washed in TBST and targets were detected using secondary antibodies conjugated to AF680 or IRDye800 diluted in Odyssey buffer. Membranes were scanned and analyzed on the Odyssey imaging system (LI-COR).

### Whole-cell lysate crosslinking

To assess OLIG2 dimerization in intact cells, whole-cell crosslinking was performed with disuccinimidyl suberate (DSS; No-Weigh format, #A39267, Thermo Fisher Scientific) reconstituted to 50 mM in anhydrous DMSO immediately before use. Approximately 5–10 × 10⁶ cells were harvested, washed three times in ice-cold PBS (pH 8.0) to remove amine-containing medium, and resuspended in 200 µL ice-cold PBS (pH 8.0). DSS was added to a final concentration of approximately 4.5 mM, and cells were incubated for 30 min at 4°C. The reaction was quenched with Tris-buffered saline (pH 7.5) for 15 min at room temperature, after which cells were washed, pelleted, and lysed as above. Crosslinked lysates were resolved by SDS-PAGE under non-reducing conditions (LDS sample buffer without reducing agent, heated at 65°C for 10 min) and immunoblotted for OLIG2, with actin as a loading control, as described above.

### Immunofluorescence

Cells were plated in Ibidi® µ-Slide 8-well chamber slides and treated with vehicle or 1 μM CT-179. Cells were then washed once in PBS and fixed with 4% PFA at room temperature for 15 min. Cells were quenched in TBS for 2 min before permeabilization and blocking in Odyssey TTX buffer containing 0.1% Tween-20 and 0.3% Triton X-100 for 1 h at 22°C. Cells were then incubated overnight at 4°C with rocking using α-tubulin mouse mAb conjugated to AF488, diluted 1:100 in Odyssey TTX. Cells were washed three times in TBS containing 0.1% Tween-20 and mounted in SlowFade™ Diamond Antifade Mountant with DAPI (Life Technologies). Images were captured using an API DeltaVision Deconvolution IX71 Olympus microscope, deconvolved using the SoftWorx program and processed using ImageJ software. For organoids, samples were fixed in 4% paraformaldehyde (PFA) in PBS, embedded in 2.5% low-melting-point agarose gel, and subsequently embedded in paraffin. Organoid sections (4 µm) were probed with anti-OLIG2 (Cell Signaling Technology) and a rabbit IgG isotype control (Cell Signaling Technology) in 10% goat serum with 0.2% Triton X-100 overnight at 4 °C, then stained with AF488-conjugated goat anti-rabbit secondary antibody. Sections were counterstained with DAPI and captured on an Aperio ScanScope slide scanner.

### Immunohistochemistry

After paraffin embedding, organoid samples were stained with hematoxylin and eosin (H&E). Organoid sections (4 µm) were probed with anti-cleaved caspase-3 (Biocare Medical) in 10% goat serum overnight at 4 °C and stained with Mach 1 Universal Polymer HRP. Signal was developed with 3,3’-diaminobenzidine (DAB). Sections were counterstained with hematoxylin, dehydrated, and mounted.

### Cellular Thermal Shift Assay (CETSA)

CETSAs were performed essentially as previously described ^15,16^. Dissociated RR2 cells (14 × 10⁶ per treatment) were treated with vehicle or 20 µM CT-179 in NSC SFM media in 15 mL tubes for 20 min at 37°C. The cells were centrifuged for 3 min at 300 *× g* and the resulting pellets were resuspended in 240 µL PBS containing Halt® inhibitor cocktail and 1 mM EDTA. For the CT-179 tests, an additional 20 µM CT-179 was added to the PBS. Cells were aliquoted into 40 µL lots into thin-wall PCR tubes and incubated in a BioRad thermocycler over an arbitrary temperature gradient between 40°C and 50°C for 3 min. Tubes were immediately moved into another thermal cycler set at 25°C for 3 min, and then snap-frozen in liquid nitrogen. Cells were freeze/thawed twice in liquid nitrogen and the thermal cycler set at 25°C with vortexing in between. Supernatants were clarified by centrifugation at 14,000 rpm for 30 min at 4°C. Lysates were subject to BCA assay in PBS diluent and 50 µg of total protein was run on 15-well 4-12% Bis-Tris gradient gels before transfer and Western blotting for OLIG2 and actin as stipulated previously.

### Electrophoretic Mobility Shift Assay (EMSA)

Cells (1 × 10⁷) were treated for 6 h with 1 µM CT-179 and nuclear extracts were harvested as previously described^48^. EMSAs were performed using the Odyssey® Infrared EMSA kit (LI-COR) and IRDye®-labeled human p27kip E-box probe and unlabeled competitor probe (IDT Technologies, Singapore) designed and validated for human OLIG2 binding specificity^19^. EMSA reactions were run on 4-20% TBE gels in 0.5× TBE buffer at 200 V at 4°C. The gel was scanned and analyzed on the LI-COR Odyssey imaging system.

### Cleaved caspase 3/7 detection

The detection of cleaved caspase 3/7 was performed according to the manufacturer’s instructions using the Caspase-Glo® 3/7 Assay (Promega) on cell lines treated for 24 h, 48 h or 72 h with vehicle or 1 µM CT-179. Luminescence was measured on an Optima FluoStar plate scanner to obtain raw data.

### Zebrafish models

Zebrafish strains used in these studies were previously described, including *Tg(Olig2:EGFP)*^20^ and CNS NB-*FOXR2*^35^. The *Tg(7.2Sox10:mCherry)* strain was generated using Gateway cloning (Invitrogen) by combining *p5E-sox10*, *pME-mCherry*, *p3E-pA* and *pDestISce1* using the Tol2 kit^49^. Drug dosing methods, imaging, and quantitation of tumor sizes were conducted as previously described^36^. Darkfield images of sox10-mCherry and olig2-GFP expression at 24 hpf embryonic were taken using an Olympus SZX16 microscope with an aperture of 0.3 and configured with an Olympus DP74-CU camera. CNS NB-FOXR2 tumor transplants before and after vehicle and drug treatments were taken using an Olympus Fluoview FV1200 confocal microscope with a UPlanSApo 10X/0.40 or UPlanSApo 60X/1.20W objective.

### Subcutaneous xenografts

GBM4 and GBM8 were obtained from the University of California San Diego cell bank; *MGMT* promoter methylation status was determined by pyrosequencing of CpG sites 74–78 (Caris Life Sciences) and was 95% in GBM4 and 55% in GBM8. Both lines were classified as MGMT-methylated by the testing laboratory; the specific methylation cutoff applied at the time is no longer available from the original report, and the GBM8 value of 55% is best regarded as borderline. For the GBM4 and GBM8 models, tumor growth was initiated on day 1 in six-week-old female NSG (NOD.Cg-*Prkdc^scid^ Il2rg^tm1Wjl^*/SzJ) mice (The Jackson Laboratory) by subcutaneous injection of 1 × 10⁷ GBM4 or GBM8 cells, respectively, into the right flank. Tumor volumes were monitored by caliper in four groups (n = 8) using the formula (width × width × length)/2. Treatments were prepared and administered beginning on day 12 post-implantation. Groups included a vehicle control cohort that received sterile water p.o. at qd × 28, a CT-179 group that received 70 mg/kg active CT-179 (105.42 mg/kg as salt) dissolved in sterile water containing 5% N,N-dimethylacetamide and 30% Captisol p.o. at qd × 28, a radiotherapy/temozolomide (RT/TMZ) group that received 1 gray (Gy) of local irradiation delivered using the Faxitron CP-160 X-ray system for 1.3 min at 130 kV and 6 mA on day 20 along with 10 mg/kg temozolomide (TMZ) p.o. on days 19 and 21, or a combination of both CT-179 and RT/TMZ treatment arms. Each animal was euthanized for tumor progression (TP) when tumor volumes reached the study endpoint of 2,000 mm^3^ or on day 152, whichever came first.

### Intracranial orthotopic xenografts

Tumor growth was initiated (day 1) in six-week-old female NSG mice by intracranial injection of 2 × 106 GBM8 cells into the right frontal cortex. Mice were randomized into two groups (n = 10) beginning on day 7 when vehicle or 20 mg/kg CT-179 dissolved in 5% N,N-dimethylacetamide and 30% Captisol with stoichiometric 0.1N hydrochloric acid was administered by intraperitoneal injection qd × 14; at survival endpoint, brain tissue was collected and stained for OLIG2 (Abcam #ab136253) and Ki-67 by Reveal Biosciences (San Diego, CA). For the oral dose-response survival study, 30 NSG mice were implanted with orthotopic GBM8 xenografts; 7 were excluded before randomization (3 died following surgery and 4 developed open surgical wounds), and the 23 surviving animals were randomized into three groups (vehicle, n = 7; 1 mg/kg, n = 8; 5 mg/kg, n = 8) and received vehicle or oral CT-179 at 1 or 5 mg/kg daily, beginning on day 7 after implantation and continued for 14 days. For the single-agent orthotopic study in adult G06, patient-derived G06 cells (3 × 105 in 3 µL) were injected intracranially into anesthetized female athymic nude (Nu/Nu) mice, which received daily oral CT-179 at 150 mg/kg/day starting on day 4 after implantation. For the GBM6 orthotopic combination study, patient-derived GBM6 cells (1 × 106) were injected intracranially into NOD-RAG mice and randomized (vehicle, n = 20; CT-179, n = 19; RT, n = 20; CT-179 + RT, n = 15) into vehicle, CT-179 (50 mg/kg intraperitoneally, three times weekly for three weeks beginning day 8), radiotherapy (2 Gy × 4 fractions over 8 days; 8 Gy total), CT-179 + RT, TMZ, TMZ + RT, or CT-179 + TMZ + RT. The TMZ-containing arms (TMZ, TMZ + RT, and CT-179 + TMZ + RT) were included to address a secondary objective and are reported separately; the present study analyzes the vehicle, CT-179, radiotherapy, and CT-179 + RT arms to isolate the contribution of CT-179 to radiotherapy in this MGMT-unmethylated model. At survival endpoint, animals were euthanized according to standard institutional veterinary guidelines for ethical animal care.

### RNA-seq of CT-179-treated glioma stem cells

Patient-derived glioma stem cell lines C419, CD133 (UT Southwestern, Bachoo laboratory) and GBM4 (University of California San Diego cell bank) were treated with vehicle (DMSO) or CT-179 at the antiproliferative IC50 dose for the indicated duration (8–12 h or 36 h; 36 libraries total, in duplicate). Total RNA was extracted using standard column-based methods, RIN equivalents (RINe) of ≥8.8 were confirmed before library preparation, and 11–41 million reads per library were generated. Reads were aligned and quantified with the TopHat / Cufflinks / Cuffmerge pipeline, and differential expression was assessed at FDR < 0.05. Gene-set enrichment analysis was performed using GSEA, and β-tubulin / OLIG2 co-expression analysis used Spearman correlation against TCGA glioblastoma RNA-seq data. The raw and processed RNA-sequencing data have been deposited in the NCBI Gene Expression Omnibus under accession GSE336473.

### Receptor / channel / transporter selectivity panel

CT-179 (1.0 × 10−5 M) was profiled across a 110-target binding and enzyme/uptake panel by Eurofins Cerep (Study #100022981). Inhibition or stimulation greater than 50% of control was considered a significant effect.

### Pharmacokinetics and brain accumulation of CT-179

Pharmacokinetic and brain-accumulation studies were conducted at BioDuro (Beijing, China). CT-179 was prepared fresh daily in a vehicle formulation of 5% N,N-dimethylacetamide and 30% Captisol (sulfobutylether-β-cyclodextrin, SBECD) in water containing 0.1N hydrochloric acid solution. In the single-dose bioavailability study, 24 C57BL/6J mice (VT River) were divided into two groups (n = 12 each) and received either a single 1 mg/kg intravenous dose or a single 20 mg/kg oral gavage dose, with three animals per timepoint. Blood samples (30 µL) were obtained by saphenous-vein puncture at 1, 4, 8, and 24 h post-dose; brain tissue was obtained at 1 h and 4 h post-dose from designated animals following euthanasia. In the steady-state CNS exposure study, 36 C57BL/6J mice were divided into 12 groups of three animals each and received once-daily oral gavage doses of 1 or 5 mg/kg for one or three consecutive days; blood samples were obtained at 1, 4, 8, and 24 h after the day-1 or day-3 dose, and brain tissue was obtained at 24 h or 48 h after the final dose. Plasma (10 µL) was precipitated using acetonitrile, and brain tissue was homogenized and clarified to a final sample volume of 20 µL. Samples were compared to an internal standard of terfenadine in acetonitrile (10 mg/mL) and blank plasma or brain samples prepared from animals not exposed to CT-179. Analysis was performed by HPLC/MS on 2 µL of plasma or 5 µL of brain sample using a Kinetex 2.6 µm C18 100Å column (30 × 3.0 mm). The mobile phases consisted of water containing 0.1% formic acid (solvent A) and 0.1% formic acid in acetonitrile (solvent B).

In a separate intraperitoneal brain-accumulation study, 18 C57BL/6J mice (VT River) were divided into six groups of three animals each and received a single intraperitoneal dose of CT-179 at 1.6 or 10 mg/kg, formulated in the same 5% N,N-dimethylacetamide and 30% Captisol vehicle and administered at 10 mL/kg (the 1.6 mg/kg level reflects the analytically verified concentration of an intended 1 mg/kg dose). Plasma was collected by saphenous-vein puncture at 0.5, 1, 4, 8, and 24 h post-dose, and brain tissue was collected at 1, 4, and 24 h after euthanasia and homogenized, with brain samples diluted six-fold and back-calculated to ng/g. Plasma and brain concentrations were determined by HPLC-MS/MS against a terfenadine internal standard, with a lower limit of quantitation of 1 ng/mL in both matrices (Supplementary Figure S6).

### Thermal Shift assays

For differential scanning fluorimetry (DSF), a truncated OLIG2 construct (“CT6”) was prepared containing the bHLH domain of OLIG2 fused to an N-terminal hexa-His tag: MGSSHHHHHHSSGLVPRGSHMTTENLYFQGAGTRLKINSRERKRMHDLNIAMDGLREVMPYAHGPSVRKLSKIATLLLARNYILMLTNSLEEMKRLVS. The MycN E-box containing dsDNA was generated by annealing two oligomers: 5’AGCTAGAAGACAGCTGTTGGAAG 3’ and 3’ TCTTCTGTCGACAACCTTCTCGA 5’ to yield a final dsDNA concentration of 358 µM. Each assay tube contained 500 µL of 1 mg/mL CT6 and either assay buffer or MycN dsDNA at a 5:1 molar ratio of dsDNA:CT6. Assay buffer was 50 mM Bis-Tris propane (pH 9.0), 500 mM arginine, and 500 mM urea. After 2 h at 22°C, 10 µM CT-179 or buffer was added, and tubes were incubated at 4°C overnight. For DSF analysis, SYPRO Orange was added at a 10:1 molar ratio to CT6 and 20 µL of sample was loaded into a PCR well. Each sample was run in duplicate. The thermal shift melting temperature was analyzed using a BioRad CFX96 Real-Time PCR machine equipped with a C1000 thermal cycler. Temperature was sampled in 0.2°C increments from 25°C to 90°C.

### Extreme limiting dilution assay

GBM4 and GBM8 cells were dissociated, plated at 3,000 to 10,000 cells per well in an ultra-low-attachment 96-well plate and incubated for 24 h at 37°C in Neurocult NS-A basal medium (#05751, StemCell Technologies, Cambridge, MA, USA) supplemented with 10 ng/mL EGF, 15 ng/mL bFGF, and 0.0002% heparin. CT-179 was added in 0.5% final DMSO at concentrations of 0, 80, 250, or 750 nM (GBM4) or 0, 50, 75, or 100 nM (GBM8), and cultures were incubated for a further 72 h. GBM spheres with a diameter greater than 80 µm were counted. Wells were counted as positive if one or more GBM spheres were present. Data were analyzed using the Extreme Limiting Dilution Assay software package (Walter and Eliza Hall Bioinformatics Institute, Melbourne, Australia).

### Real-time reverse-transcription quantitative PCR

All cells were treated with either vehicle or 1 μM of CT-179 for 24 h. RNA was extracted using the RNeasy® Mini Kit (QIAGEN), with cDNA synthesis conducted on 2–5 µg of RNA as per the manufacturer’s instructions using the SuperScript® III First-Strand synthesis supermix kit (Invitrogen). Quantitative PCR was performed on the QuantStudio 6 Flex Real-Time PCR machine (Life Technologies) as previously described^23^ using pre-optimized FAM-labeled primer-probe mixes for *Olig2*, *Cdk1*, *Cdk2*, *Cdk4*, *Cdk6*, *Rb*, *Pdgfr-α*, *Met*, *Egfr* and *Erbb4* and VIC-labeled primer-limited primer-probe mix for *Actb* as an endogenous control (all supplied by Life Technologies). For the organoid and QBC512 analyses, total RNA was extracted using the RNeasy Mini Kit (Qiagen) according to the manufacturer’s instructions, first-strand cDNA was synthesized using SuperScript IV (Invitrogen), and relative quantitation was performed with SYBR Green PCR Master Mix (Applied Biosystems) in triplicate on an ABI ViiA 7 instrument (Applied Biosystems). Cycle thresholds were determined and exported using QuantStudio software (Applied Biosystems), and transcript levels were determined relative to the β-actin (ACTB) reference gene.

### Pulse-chase labeling

Pulse labeling of nascent proteins and subsequent analysis were conducted as previously described^23^.

### Statistical Analysis

Data are presented as mean ± standard error of the mean (s.e.m.) unless otherwise indicated. Group comparisons used unpaired two-tailed Student’s t-test for two-group analyses (Fig. 1g, OPC quantitation; Fig. 4f and Fig. 4e endpoint comparisons of combination versus RT/TMZ alone) and one-way ANOVA for three or more groups (Fig. 2c, cleaved caspase 3/7; Fig. 4b, zebrafish tumor area; Fig. 4e and Fig. 4f, day-50 and day-42 multi-arm comparisons; Fig. 4e and Fig. 4f, day-74 and day-61 RT/TMZ-versus-combination comparisons). Survival differences were assessed by log-rank (Mantel-Cox) test (Fig. 4d, Supplementary Figures S2 and S4, and Fig. 5). Tubulin / OLIG2 expression correlations in TCGA glioblastoma specimens were assessed by Spearman rank correlation with FDR adjustment. Significance is denoted as * *P* < 0.05, ** *P* < 0.01, and *** *P* < 0.001; exact p values are reported in figure legends where space permits. Half-maximal inhibitory concentrations (IC50) and stem-cell frequency estimates were calculated using nonlinear regression and the Extreme Limiting Dilution Assay software (Walter and Eliza Hall Bioinformatics Institute), respectively. For the organoid experiments, a polygon selection tool in ImageJ was used to outline organoids on brightfield images before and after treatment; measurement of the outlines yielded two-dimensional organoid area and a circularity measurement, reported here as sphericity. Slide-scanner images were imported into QuPath, stain vectors were analyzed and adjusted, and positive-cell detection was used to quantify positive-cell populations, with cleaved caspase-3-positive cells counted computationally and manually verified. Whole organoids were treated as individual biological replicates; morphometric analyses were performed on six organoids per condition and staining quantifications on three to six organoids per condition, as some organoids were lost during paraffin embedding owing to their small size. Organoid comparisons were assessed by two-way ANOVA. All analyses were performed in GraphPad Prism (version 9 or later); n values for each experiment are stated in the corresponding figure legend.

---

**Supplementary Figure S1:**
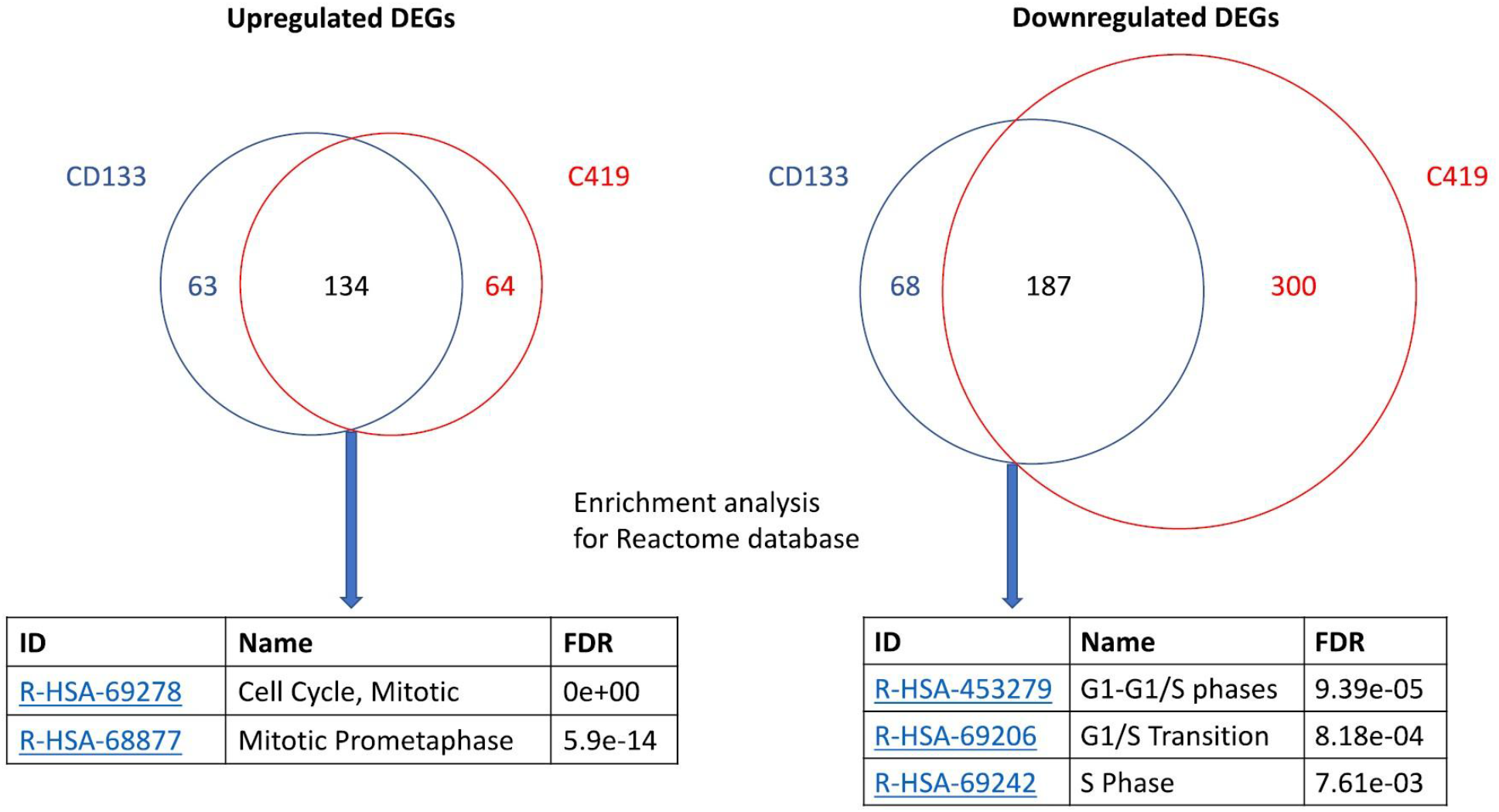
Late-time-point gene-expression signature of CT-179-treated glioma stem cells. Venn diagrams of differentially expressed genes in C419 and CD133 patient-derived glioma stem cells at 36 h after CT-179 treatment (FDR < 0.05 and fold change > 1.5 or < −1.5), showing 134 concordantly up-regulated and 187 concordantly down-regulated genes shared between the two lines. Tables show the most enriched Reactome pathways for the shared gene sets. Shared up-regulated genes are enriched for Cell Cycle, Mitotic (FDR 0e+00) and Mitotic Prometaphase (FDR 5.9e-14); shared down-regulated genes are enriched for G1/G1-S phases (FDR 9.39e-05), G1/S Transition (FDR 8.18e-04) and S phase (FDR 7.61e-03), consistent with cells arresting in mitosis. Source: UT Southwestern (Kittler laboratory).

**Supplementary Figure S2:**
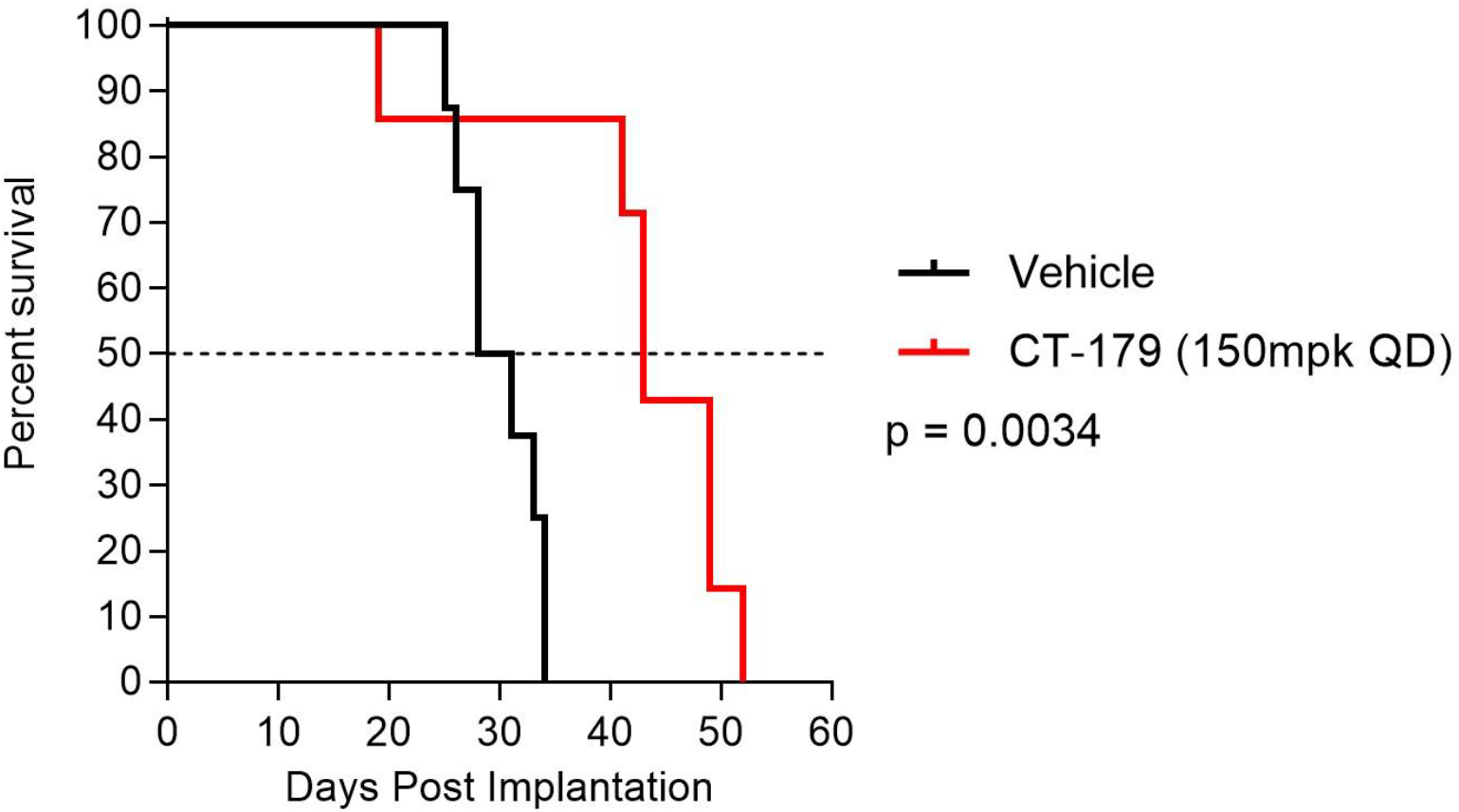
Single-agent CT-179 in an additional adult orthotopic background. Kaplan-Meier survival curves for athymic nude mice (Nu/Nu) bearing intracranial G06 patient-derived xenografts (3 × 10⁵ cells per implant; n = 7 per arm) randomized to vehicle or oral CT-179 (150 mg/kg/day, initiated on Day 4). Median survival was 29.5 days for vehicle-treated and 43.0 days for CT-179-treated animals, an approximately 46% extension (*P* = 0.0034, log-rank Mantel-Cox test). Body weights remained comparable to vehicle controls throughout the study.

**Supplementary Figure S3:**
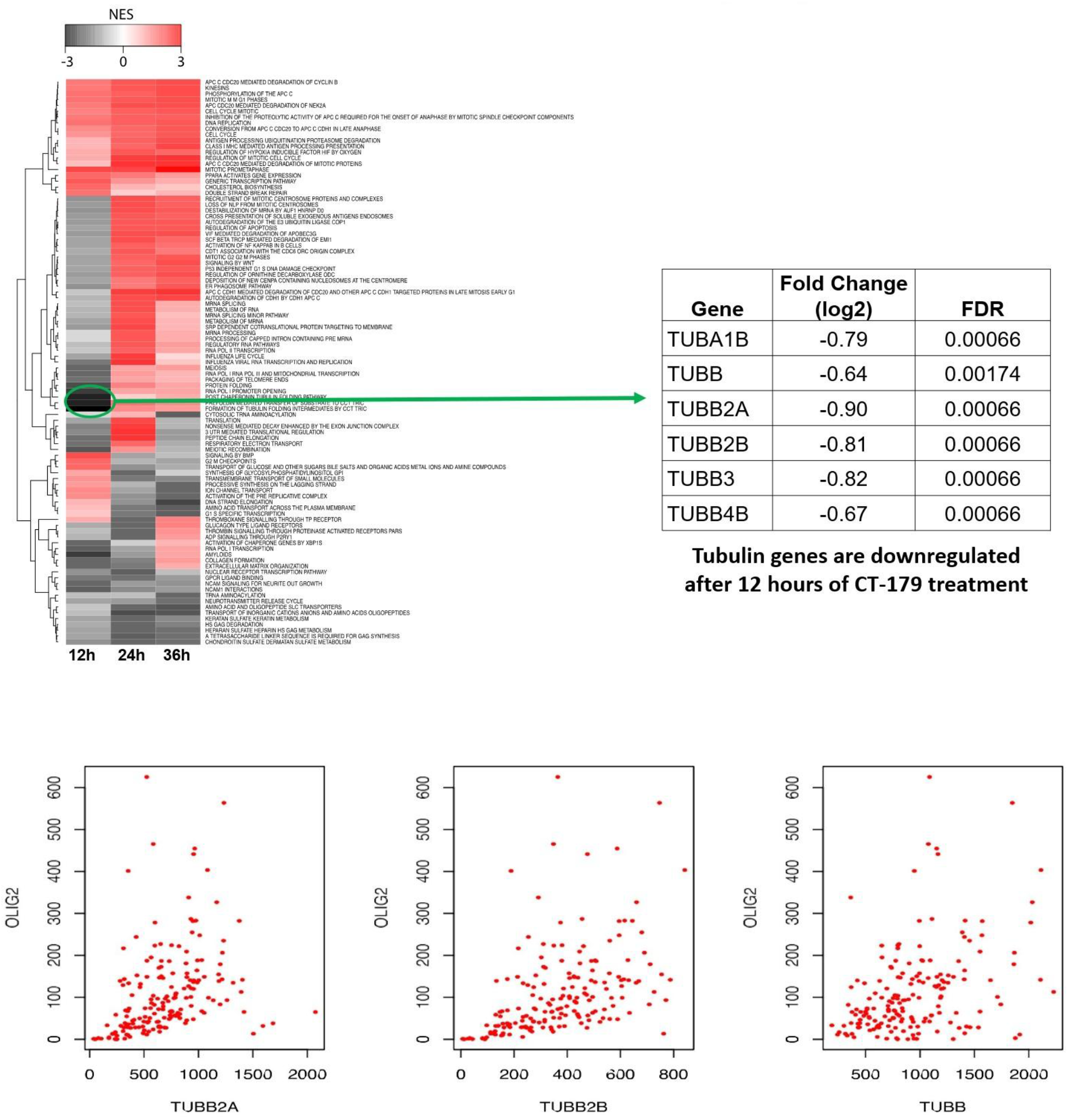
CT-179 represses the β-tubulin transcriptional program in patient-derived glioma stem cells. RNA-seq analysis of CT-179-treated patient-derived glioma stem cell lines (C419, CD133, GBM4) at early (8–12 h) and late (36 h) time points (36 libraries; FDR < 0.05). Top: hierarchical-clustering heat map of GSEA normalized enrichment scores across CT-179 treatment time points (12, 24, and 36 h) in C419 cells, with the post-chaperonin tubulin-folding pathway cluster highlighted; an accompanying table reports log₂ fold change and FDR for the most strongly down-regulated β-tubulin transcripts at 12 h (*TUBA1B*, −0.79; *TUBB*, −0.64; *TUBB2A*, −0.90; *TUBB2B*, −0.81; *TUBB3*, −0.82; *TUBB4B*, −0.67; FDR ≤ 0.00174 for all). Bottom: Spearman correlation of *TUBB2A*, *TUBB2B*, and *TUBB* expression with OLIG2 expression in TCGA glioblastoma specimens (ρ = 0.551, 0.579, and 0.401, respectively; FDR-significant), nominating these β-tubulin isoforms as putative direct OLIG2 transcriptional targets in patient tumors, although direct OLIG2 promoter occupancy at these loci was not assessed. The full late-timepoint differential-expression analysis, demonstrating concordant up-regulation of G2/M-phase transcripts and down-regulation of G1/S-phase transcripts at 36 h consistent with cells arresting in mitosis, is provided in Supplementary Figure S1.

**Supplementary Figure S4:**
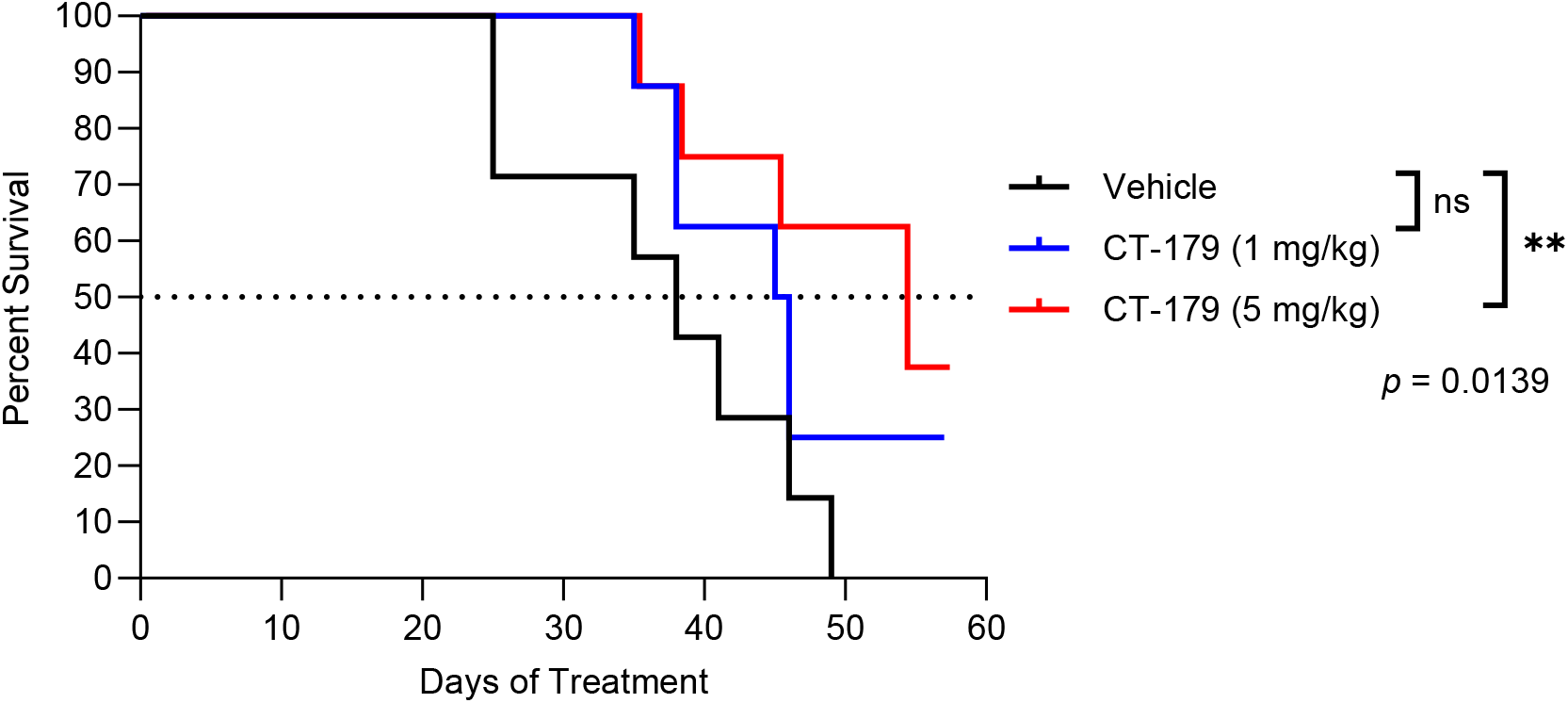
Oral CT-179 produces a dose-dependent extension of survival in orthotopic GBM8 xenografts. Kaplan-Meier survival curves for NSG mice bearing intracranial GBM8 patient-derived xenografts treated with vehicle or oral CT-179 at 1 or 5 mg/kg daily until sacrificed (Control, n = 7; CT-179 1 mg/kg, n = 8; CT-179 5 mg/kg, n = 8). Daily oral CT-179 at 5 mg/kg conferred an approximately 42% extension in median survival relative to vehicle by log-rank (Mantel-Cox) test (*P* = 0.0139), whereas the 1 mg/kg arm did not reach significance.

**Supplementary Figure S5:**
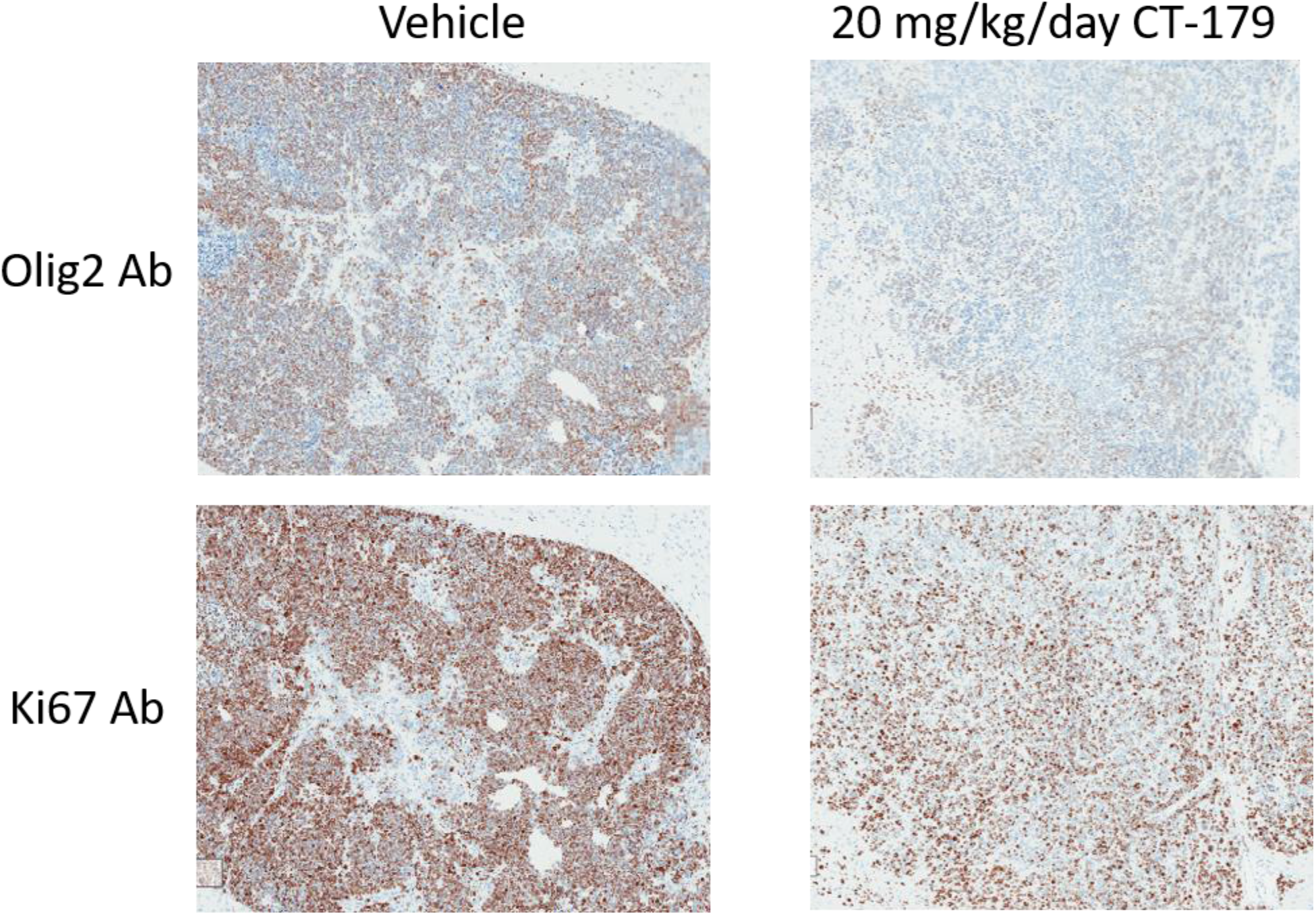
CT-179 is associated with reduced intratumoral OLIG2 and Ki-67 staining in orthotopic GBM8 xenografts. Representative immunohistochemistry of resected tumor tissue from vehicle- and CT-179-treated NSG mice (CT-179 20 mg/kg/day intraperitoneally for 14 days, beginning day 7 post-implantation) stained for OLIG2 or Ki-67 (brown) and counterstained with hematoxylin (blue). Representative panels suggest reductions in OLIG2 and Ki-67 staining in CT-179-treated tumors, consistent with in vivo target engagement and reduced tumor cell proliferation. These are qualitative, representative images, and staining was not quantified.

**Supplementary Figure S6:**
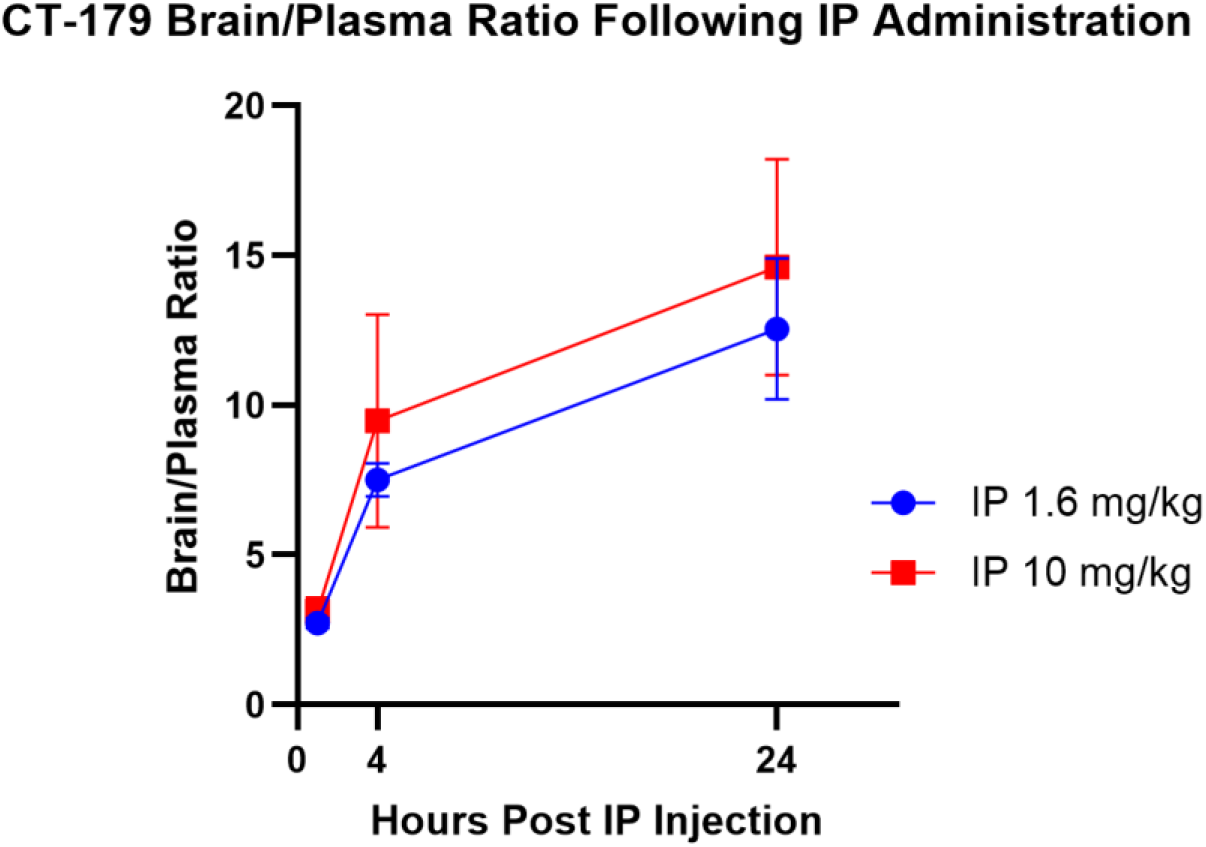
CT-179 achieves comparable brain exposure following oral and intraperitoneal administration. Brain and plasma concentrations of CT-179 in C57BL/6J mice following intraperitoneal administration at 1.6 or 10 mg/kg, sampled at 1, 4, and 24 h. Brain-to-plasma ratios increased over time to approximately 10–15 at 24 h at both doses, comparable to the brain-to-plasma ratios observed after oral dosing (Fig. 4c), indicating that CNS exposure of CT-179 is similar across the oral and intraperitoneal routes.

**Supplementary Figure S7:**
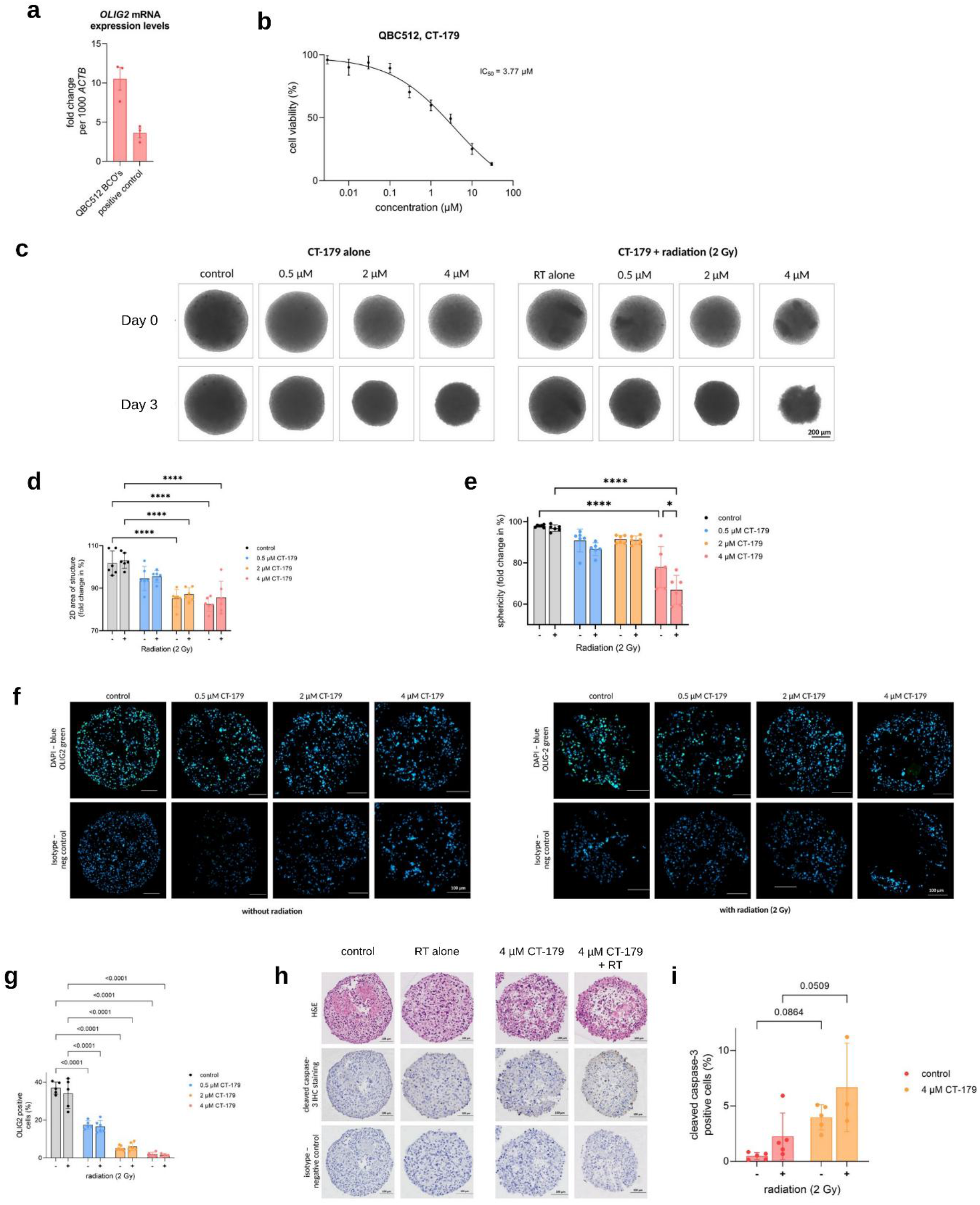
CT-179 dose-dependently depletes OLIG2, compromises organoid integrity, and induces apoptosis in patient-derived GBM organoids. Glioblastoma organoids derived from freshly resected patient tumor tissue (line QBC512) were treated with 0.5, 2, or 4 µM CT-179 with or without a single 2 Gy radiotherapy (RT) fraction and analyzed at 72 h. a. OLIG2 mRNA expression in QBC512 organoids relative to the OLIG2-positive DAOY medulloblastoma line (positive control) by RT-qPCR, normalized to ACTB (QBC512 relative expression values 12, 11, and 7; n = 3 technical replicates). b. CT-179 dose-response in the matched adherent QBC512 line by sulforhodamine B assay at 72 h; IC50 = 3.77 µM (n = 3 technical replicates). c. Representative brightfield images at day 0 and day 3 across the CT-179 dose range with and without RT (scale bar, 200 µm). d. Organoid 2D area and e. sphericity, expressed as fold change from baseline (n = 6 organoids, each treated as an individual biological replicate). f. Representative OLIG2 immunofluorescence across the dose range with and without RT (OLIG2, green; DAPI, blue), with isotype-control panels (scale bar, 100 µm). g. Quantification of OLIG2-positive cells (n ≥ 3 organoids per condition; each dot represents one organoid). h. Representative hematoxylin and eosin and cleaved caspase-3 immunohistochemistry (control, RT alone, 4 µM CT-179, and 4 µM CT-179 + RT), with isotype-control panels (scale bar, 100 µm). i. Quantification of cleaved caspase-3-positive cells in control and 4 µM CT-179 organoids with and without RT (n ≥ 3 organoids per condition; each dot represents one organoid). Each organoid was treated as an individual biological replicate, and for each organoid two adjacent sections were stained, one for the target antibody and one for the isotype (negative) control. Data are from a single experiment and are presented as mean ± SD or SEM as indicated; P values were determined by two-way ANOVA.

## Notes

https://www.ncbi.nlm.nih.gov/geo/

